# Shiny-Calorie: A context-aware application for indirect calorimetry data analysis and visualization using R

**DOI:** 10.1101/2025.04.24.648116

**Authors:** Stephan Grein, Tabea Elschner, Ronja Kardinal, Johanna Bruder, Akim Stromeyer, Karthikeyan Gunasekaran, Jennifer Witt, Hildigunnur Hermannsdóttir, Janina Behrens, Mueez U-Din, Jiangyan Yu, Gerhard Heldmaier, Renate Schreiber, Jan Rozman, Markus Heine, Ludger Scheja, Anna Worthmann, Jörg Heeren, Dagmar Wachten, Kerstin Wilhelm-Jüngling, Alexander Pfeifer, Jan Hasenauer, Martin Klingenspor

## Abstract

Indirect calorimetry is a cornerstone technique for metabolic phenotyping of animal models in preclinical research, with well-established experimental protocols and platforms. However, a flexible, extensible, and user-friendly software suite that enables standardized integration of data and metadata from diverse metabolic phenotyping platforms—followed by unified statistical analysis and visualization—remains absent. We present Shiny-Calorie, an open-source interactive web application for transparent data and metadata integration, comprehensive statistical data analysis, and visualization of indirect calorimetry datasets. Shiny-Calorie is compatible with data formats from widely used commercial metabolic phenotyping platforms, such as TSE and Sable Systems, and includes functionality for exporting processed data in these formats. Built using GNU R and a Shiny-based reactive interface, Shiny-Calorie enables intuitive exploration of complex, multi-modal longitudinal datasets comprising categorical, continuous, ordinal, and count variables. The platform incorporates state-of-the-art statistical methods for robust hypothesis testing, thereby facilitating biologically meaningful interpretation of energy metabolism phenotypes, including resting metabolic rate and energy expenditure. Overall, Shiny-Calorie streamlines routine analysis workflows and enhances reproducibility and transparency in metabolic phenotyping studies.

## 1 Introduction

Indirect calorimetry (IC) is an indispensable tool for metabolic phenotyping [1], and is routinely employed in pre-clinical studies and clinical research, particularly in the domains of adipose tissue research [2, 3, 4, 5] and nutritional medicine [6]. IC is used to quantify both variations and mean differences in energy expenditure by tracking the respiratory gases – oxygen and carbon dioxide. These measurements enable the analysis of correlations and effect sizes related to genotype or diet stratifications, typically corrected for variations in body composition. IC has been utilized to identify and characterize metabolic phenotypes for decades [7, 8, 9, 10, 11]. It also allows for the calculation of resting metabolic rates, which are especially relevant in the context of phenotyping in obesity research [2, 6, 12].

Several commercial and academic software packages exist for the processing and visualization of IC data, such as the proprietary analysis tools provided by TSE and Sable Systems, as well as standalone packages tailored to specific institutional workflows [13, 14]. In addition, various R and Python scripts are available within the research community; however, these scripts often lack generalizability, user-friendly interfaces, and support for harmonized metadata integration. Existing solutions typically focus on individual experiments, and poorly support cross-study analyses, and offer limited extensibility or reproducibility. In most cases, metadata must be manually curated, and labels are not harmonized across datasets, complicating efforts to conduct joint analyses across cohorts or institutions. Additionally, there are no open-source platforms that provide graphical user interfaces and support multi-modal longitudinal datasets.

In this study we present the Shiny-Calorie web application with its streamlined user interface for seamless integration of data and harmonization of metadata. Shiny-Calorie thus enables context-aware data visualization and analysis. The application was developed in collaboration with experienced IC researchers and is tailored to their requirements for data processing, analysis, and visualization. Accordingly, Shiny-Calorie alleviates the above mentioned issues by providing a unified and customizable analysis workflow which also automatically integrates data with metadata. Shiny-Calorie enables the quantification of energy expenditure (EE), resting metabolic rate (RMR), and activity-dependent energy expenditure (AEE) and the linking of the results with information about genotype, food and water intake, physical activity, etc. (Fig. 1, Supplementary Figures S1-S3). Implemented in the GNU R ecosystem, Shiny-Calorie provides access to state-of-the-art visualization and statistical analysis tools. Publication-ready high-quality figures and standardized data export [13] of computed and consolidated datasets for downstream analysis are provided. Shiny-Calorie is available as (i) a platform-independent web application running in common web browsers (applicable when users lack administrator privileges or do not want to install software), (ii) a standalone Desktop application (applicable when connectivity is limited), and (iii) a Docker container (promoting a quick adoption by users and instant transferability of workflows between workstations).

**Figure 1.**
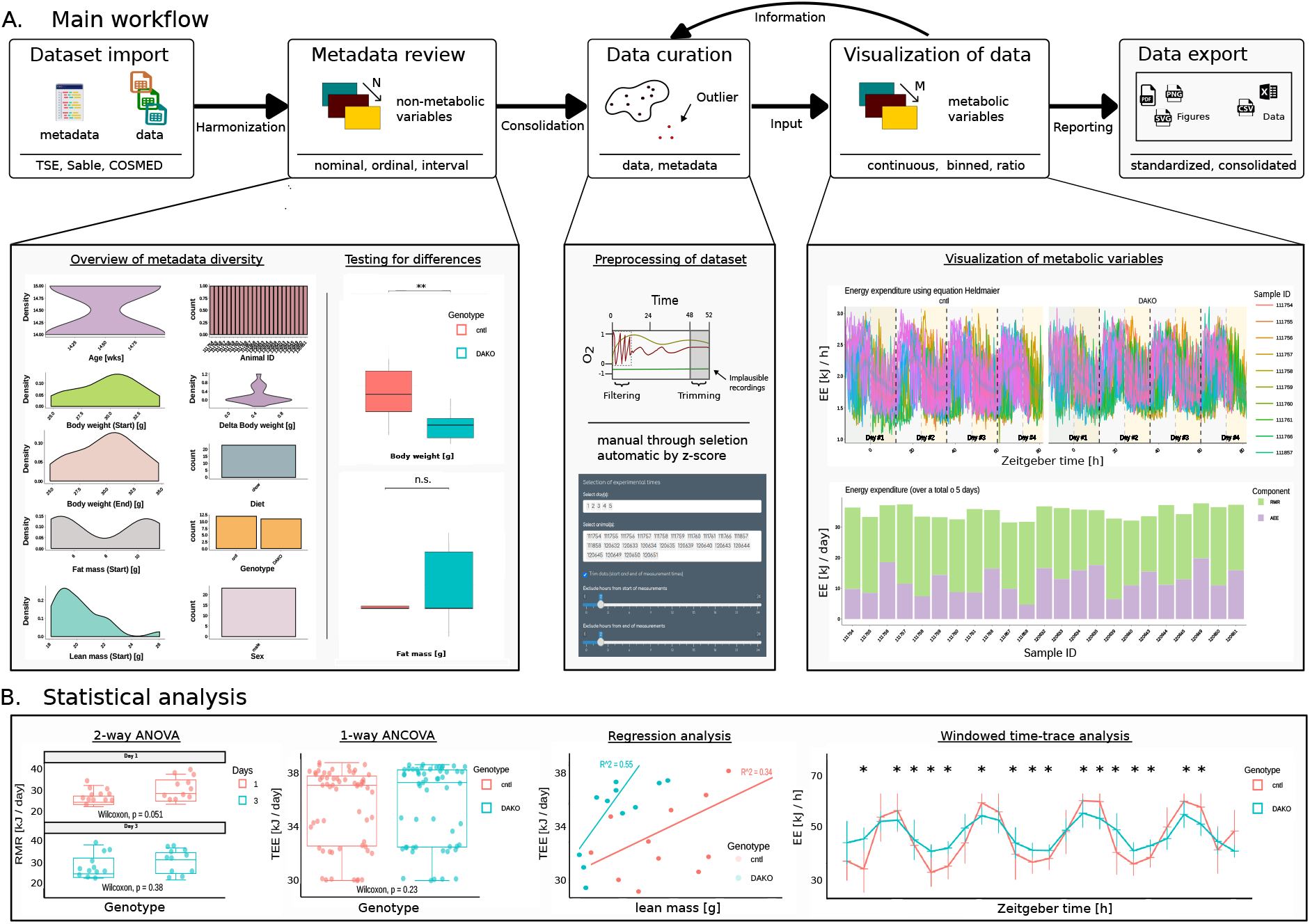
Shiny-Calorie workflow and features. (A) Illustration of the Shiny-Calorie workflow: dataset import, metadata review, data curation, visualization of data and data export. Exemplary visualizations for different datasets are shown, i.e. bar, density, box, silhouette and time-series plots. (B) Overview of statistical analysis methods and corresponding visualization, including statistical analysis with ANOVA and ANCOVA, regression analysis and time-series analysis.

## 2 Feature Overview

Shiny-Calorie offers a comprehensive set of features for the integration, processing, analysis, and visualization of indirect calorimetry (IC) data. The following sections provide a detailed description of the software’s key capabilities, organized according to the typical user workflow. Advanced features and extended options are illustrated in the Supporting Information (SI) and demonstrated through four representative use-cases.

### 2.1 Dataset Import

Shiny-Calorie enables the import of datasets from multiple metabolic phenotyping platforms, including TSE Systems, Sable Systems, CaloBox and COSMED. Supported file formats and versions are listed in Supplemental Table S1. At a minimum, imported datasets must contain an `Animal ID` column and metabolic variables such as *O*_2_ and *CO*_2_, typically recorded at regular time intervals. Shiny-Calorie accepts raw exports directly from metabolic phenotyping platforms and can ingest data files containing metadata either in headers or in structured external files. Moreover, Shiny-Calorie also supports ontology-based hierarchical metadata via a standardized Metadata sheet [15] for enhanced data annotation and interoperability.

### 2.2 Metadata Review

Metadata in Shiny-Calorie can be supplied in two ways: either manually, through user input during the data review process, by uploading a standardized Metadata sheet [15], or through an auxiliary Shiny app assisting with metadata collection. The Metadata sheet enables structured annotation of datasets with experimental conditions, treatments, photoperiod, and other contextual variables. Even if metadata are imported, the user can manually change the metadata, e.g., to correct errors or to fill gaps.

Once metadata is provided—either via manual input or upload—Shiny-Calorie performs automatic harmonization of metadata labels across datasets. This includes consolidating variable names, resolving inconsistencies, and encoding the metadata into a standardized internal format to support joint analysis across multi-cohort studies. Harmonization reduces the risk of errors from inconsistent labeling and facilitates reproducible downstream analysis.

### 2.3 Data Curation

Shiny-Calorie provides robust tools for the cleaning and curation of indirect calorimetry datasets to ensure data quality and consistency prior to downstream analysis. To ensure data integrity, users may enable a set of pre-defined consistency checks via checkboxes during the metadata review step. These checks include the identification of negative values in respiratory gas measurements, filtering of high-frequency data points, enforcement of complete measurement days, and validation of temperature records.

Upon data upload, these consistency checks are automatically applied to detect common anomalies in the raw measurements. Users are notified of issues and can decide on appropriate corrective actions. Outliers can be interactively removed using graphical tools, including rectangular and lasso selection, directly within the plotting interface, or through adjustable z-score thresholding.

Individual subjects and days can be excluded from the dataset based on metadata attributes or statistical outlier detection. Additionally, grouping and filtering based on any categorical metadata variable (e.g., genotype, treatment, diet) are supported to facilitate stratified and comparative analyses.

To eliminate artifacts introduced during the initial habituation phase or final handling of subjects, users can trim time segments from the beginning and end of experiments. Reliable measurement periods, such as intermediate full days, can be selected using dropdown menus. Selection can be performed either based on zeitgeber time (aligned to light-phase onset) or conventional calendar dates. Photoperiod information, if not present in the raw data, is inferred from the Metadata sheet or specified manually.

### 2.4 Data Processing and Visualization

With harmonized datasets, Shiny-Calorie reconstructs energy expenditure (EE) from raw respiratory gas exchange measurements using standard equations, including the Heldmaier equation [16],

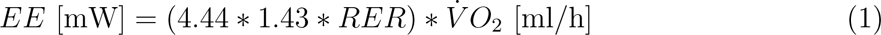

where 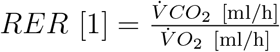.

Alternative equations are also supported (cf. Supplemental Table S2). Shiny-Calorie enables derivation of secondary quantities such as RMR and activity-dependent energy expenditure (AEE), using methods based on the variability of *O*_2_/*CO*_2_ signals (see SI Methods). When physical activity data is available, Shiny-Calorie generates locomotion density maps and behavioral summaries for experimental validation.

Derived metrics are calculated to assess substrate utilization (e.g., fat vs. carbohydrate oxidation). Visualizations of raw and reference *O*_2_/*CO*_2_ signals assist in evaluating data quality and validating experimental conditions. Shiny-Calorie supports the plotting of all available variables—metabolic and non-metabolic—such as food/water intake, locomotor activity and photoperiods. Windowed time-trace analyses allow for exploration of temporal dynamics (e.g., hourly RMR/EE profiles), enabling detection of time-specific differences between groups that may be masked by averaging.

### 2.5 Statistical Analysis

Shiny-Calorie integrates standard statistical methods for hypothesis testing. Available models include multi-way ANOVA, ANCOVA, and generalized linear models, with support for variable number of comparison groups. Covariate correction (e.g., for body weight) is available [17]. Diagnostic tools and assumption tests (e.g., Shapiro–Wilk, Levene’s test) are provided to verify prerequisites for parametric testing. When assumptions are violated, Shiny-Calorie either automatically switches to non-parametric alternatives or allows the user to select them manually. A decision-support flowchart for statistical testing is provided in the SI.

Post-hoc comparisons (with correction for multiple testing) and statistical summaries are generated, and test assumption results are summarized in a binary format (check marks), facilitating interpretation for users without statistical expertise. Statistical significance is reported using conventional asterisk notation.

### 2.6 Data Export

Users can export all processed and derived data in multiple formats. Visualizations are available in vector-based (SVG/PDF), bitmap formats (PNG) and as interactive plots (HTML). Data tables can be exported as CSV files, while consolidated multi-cohort datasets can be saved in Excel or CalR-compatible formats [13]. These exports enable downstream analysis, facilitate reproducibility, and allow cross-study comparisons. The `Export` tab allows users to download merged data frames suitable for offline exploration and integration into other computational workflows.

## 3 Dissemination, Training and Use-cases

To support dissemination, training, and user adoption, Shiny-Calorie is accompanied by a comprehensive suite of instructional resources and real-world examples.

### Documentation

Extensive documentation of Shiny-Calorie’s functionalities, including API references and deployment instructions, is available on the project’s documentation page: https://ICB-DCM.github.io/Shiny-Calorie

### Use-cases

Four representative use-cases are provided in the Supporting Information (SI) to demonstrate Shiny-Calorie’s capabilities in real-world scenarios.

- The introductory use-case illustrates how to streamline metadata harmonization using the standardized Metadata sheet [15], avoiding laborious manual annotation tasks.
- The second use-case discusses two example IC datasets (DAKO and UCP1-KO studies) to showcase step-by-step the application of Shiny-Calorie’s main features, i.e. preprocessing, visualization, and statistical data analysis
- The third use-case demonstrates the possibility to analyze additionally the locomotor activity and feeding/drinking patterns of subjects (and thus how to exploit IC datasets comprehensively)
- The last use-case shows one of Shiny-Calorie’s advanced statistical features, i.e. to employ wavelet analysis to study ultradian rhythms in Djungarian hamsters (Supplementary Figures S4-S5)
- These datasets are bundled with the application and can be directly loaded using the Load example data buttons on the landing page. This enables users, including those without their own IC datasets, to explore Shiny-Calorie’s full functionality.

### Video Materials

Instructional screen recordings are available on YouTube: http://youtube.com/@Shiny-Calorie The web application also includes an interactive inapp guide (accessible via the Guide button) and context-specific help sections (accessible via the Help button). These resources are designed to facilitate onboarding and self-paced training.

### Training-Oriented Design

Shiny-Calorie’s analysis workflows were developed in close collaboration with experimentalists to optimize usability for routine laboratory applications. Design decisions prioritize transparency, reproducibility, and ease of use, ensuring that even non-specialist users can efficiently perform and interpret metabolic analyses.

## 4 Implementation and Availability

Shiny-Calorie is implemented as a web-based application using the `Shiny` framework within the GNU R ecosystem. The interface is reactive and modular, enabling dynamic interaction and real-time data exploration. Visualization is built upon the `Plotly` library [18], facilitating high-quality, interactive plotting of multi-dimensional longitudinal datasets.

The application is freely available under the BSD-3-Clause license. Shiny-Calorie can be accessed in three deployment modes:

- **Web Application:** Hosted on institutional infrastructure and accessible via standard web browsers without the need for installation^1^.
- **Docker Image:** Available as an OCI-compliant container for rapid deployment and reproducible analysis workflows^2^.
- **Standalone Desktop Installer:** Provided via Github for offline use on local machines without requiring internet access^3^.

The source code and API references are available on GitHub^4^. Feature requests, bug reports, and contributions are welcomed via the GitHub issue tracker and pull request system.

## 5 Discussion

Shiny-Calorie addresses a key bottleneck in the field of metabolic phenotyping by enabling harmonized, reproducible, and interactive analysis of indirect calorimetry datasets across multiple platforms and experimental contexts. Its modular and extensible architecture supports both single-study and multi-cohort analyses, enhancing the utility of IC data in preclinical and translational research.

By automating data harmonization and integrating robust statistical and visualization methods into a single platform, Shiny-Calorie substantially reduces the overhead of metadata curation and manual processing. It supports stratified analysis of metabolic parameters (e.g., EE, RMR, AEE) with or without physical activity data, making it adaptable to a wide range of experimental designs and instrumentation configurations.

Shiny-Calorie’s design reflects the practical needs of both computational analysts and wetlab researchers. Through an intuitive graphical interface and guided workflows, it empowers users with varying levels of technical expertise to perform high-quality metabolic data analyses. The platform’s open-source nature and multi-platform availability foster transparency, reproducibility, and collaborative development.

Overall, Shiny-Calorie contributes a broadly applicable and user-centered solution to the metabolic research community, supporting standardized, scalable, and insightful interrogation of energy metabolism in diverse biological systems.

## Supporting information

Shiny-Calorie-Supplemental

## 7 Acknowledgments

The icon ‘programming-language’ by `juicy-fish` and the icons for file types by `Freepik` from `Flaticon.com` have been used to design the workflow and features figure.

## Footnotes

1 https://shiny.iaas.uni-bonn.de/Shiny-Calorie

2 https://hub.docker.com/r/stephanmg/Shiny-Calorie

3 https://github.com/ICB-DCM/Shiny-Calorie-Wrapper

4 https://github.com/ICB-DCM/Shiny-Calorie

## References

[1] Patrick C. Even and Nachiket A. Nadkarni. Indirect calorimetry in laboratory mice and rats: principles, practical considerations, interpretation and perspectives. American Journal of Physiology-Regulatory, Integrative and Comparative Physiology, 303(5): R459–R476, 2012. doi: 10.1152/ajpregu.00137.2012. URL https://doi.org/10.1152/ajpregu.00137.2012. PMID: 22718809.

[2] Sebastian Dieckmann, Akim Strohmeyer, Monja Willershaüser, Stefanie F. Maurer, Wolfgang Wurst, Susan Marschall, Martin Hrabe de Angelis, Ralf Kühn, Anna Worthmann, Marceline M. Fuh, Joerg Heeren, Nikolai Köhler, Josch K. Pauling, and Martin Klingenspor. Susceptibility to diet-induced obesity at thermoneutral conditions is independent of UCP1. American Journal of Physiology-Endocrinology and Metabolism, 322(2):E85–E100, 2022. doi: 10.1152/ajpendo.00278.2021. URL https://doi.org/10.1152/ajpendo.00278.2021. PMID: 34927460.

[3] Alexander W. Fischer, Christian Schlein, Barbara Cannon, Joerg Heeren, and Jan Nedergaard. Intact innervation is essential for diet-induced recruitment of brown adipose tissue. American Journal of Physiology-Endocrinology and Metabolism, 316(3): E487–E503, 2019. doi: 10.1152/ajpendo.00443.2018. URL https://doi.org/10.1152/ajpendo.00443.2018. PMID: 30576247.

[4] M. u Din, J. Raiko, T. Saari, N. Kudomi, T. Tolvanen, V. Oikonen, J. Teuho, H. T. Sipilä, N. Savisto, R. Parkkola, P. Nuutila, and K. A. Virtanen. Human brown adipose tissue [15O]O2 PET imaging in the presence and absence of cold stimulus. EJNMMI, 43, 2006. doi: 10.1007/s00259-016-3364-y.

[5] Katarina Klepac, JuHee Yang, Staffan Hildebrand, and Alexander Pfeifer. Rgs2: A multifunctional signaling hub that balances brown adipose tissue function and differentiation. Molecular Metabolism, 30:173–183, 2019. ISSN 2212-8778. doi: 10.1016/j.molmet.2019.09.015. URL https://www.sciencedirect.com/science/article/pii/S2212877819309184.

[6] Martin Klingenspor, Tobias Fromme, and Stefanie Maurer. Ablation of uncoupling protein 1 causes opposing changes of glucose uptake in brown and brite fat compatible with regulation of Glut 4 gene expression. The FASEB Journal, 34(S1): 1–1, 2020. doi: 10.1096/fasebj.2020.34.s1.06713. URL https://faseb.onlinelibrary.wiley.com/doi/abs/10.1096/fasebj.2020.34.s1.06713.

[7] E Ravussin, S Lillioja, T E Anderson, L Christin, and C Bogardus. Determinants of 24-hour energy expenditure in man. methods and results using a respiratory chamber. The Journal of Clinical Investigation, 78(6):1568–1578, 12 1986. doi: 10.1172/JCI112749. URL https://www.jci.org/articles/view/112749.

[8] James A. Levine, Norman L. Eberhardt, and Michael D. Jensen. Role of nonexercise activity thermogenesis in resistance to fat gain in humans. Science, 283(5399):212–214, 1999. doi: 10.1126/science.283.5399.212. URL https://www.science.org/doi/abs/10.1126/science.283.5399.212.

[9] Mikael Bjursell, Anna-Karin Gerdin, Christopher J. Lelliott, Emil Egecioglu, Anders Elmgren, Jan Törnell, Jan Oscarsson, and Mohammad Bohlooly-Y. Acutely reduced locomotor activity is a major contributor to western diet-induced obesity in mice. Am J Physiol Endocrinol Metab, 294(2):E251–E260, 2008. doi: 10.1152/ajpendo.00401.2007. URL https://doi.org/10.1152/ajpendo.00401.2007. PMID: 18029443.

[10] Roland L Weinsier, Gary R Hunter, Renée A Desmond, Nuala M Byrne, Paul A Zuckerman, and Betty E Darnell. Free-living activity energy expenditure in women successful and unsuccessful at maintaining a normal body weight1,2,3. J Clin Nutr, 75 (3):499–504, 2002. ISSN 0002-9165. doi: 10.1093/ajcn/75.3.499. URL https://www.sciencedirect.com/science/article/pii/S0002916523061476.

[11] Colleen M. Novak, Catherine M. Kotz, and James A. Levine. Central orexin sensitivity, physical activity, and obesity in diet-induced obese and diet-resistant rats. Am J Physiol, 290(2):E396–E403, 2006. doi: 10.1152/ajpendo.00293.2005. URL https://doi.org/10.1152/ajpendo.00293.2005. PMID: 16188908.

[12] Hui Wang, Monja Willershaüser, Yongguo Li, Tobias Fromme, Katharina Schnabl, Andrea Bast-Habersbrunner, Samira Ramisch, Sabine Mocek, and Martin Klingenspor. Uncoupling protein-1 expression does not protect mice from diet-induced obesity. American Journal of Physiology-Endocrinology and Metabolism, 320(2):E333–E345, 2021. doi: 10.1152/ajpendo.00285.2020. URL https://doi.org/10.1152/ajpendo.00285.2020. PMID: 33252252.

[13] Amir I. Mina, Raymond A. LeClair, Katherine B. LeClair, David E. Cohen, Louise Lantier, and Alexander S. Banks. Calr: A web-based analysis tool for indirect calorimetry experiments. Cell Metab, 28(4):656–666.e1, 2018. ISSN 1550-4131. doi: 10.1016/j.cmet.2018.06.019. URL https://www.sciencedirect.com/science/article/pii/S1550413118304017.

[14] A. Mina and T. Stalder. calr2: An r package for indirect calorimetry. bioRxiv, 2024. In preparation.

[15] L. Seep, S. Grein, I. Splichalova, et al. From Planning Stage To FAIR Data: A Practical Metadatasheet For Biomedical Scientists. Sci Data, 11, 2024.

[16] G. Heldmaier. Metabolic and thermoregulatory responses to heat and cold in the djungarian hamster, phodopus sungorus. J. Comp. Phys., pages 115–122, 1975.

[17] T.D Müller, M. Klingenspor, and M. H. Tschöp. Revisiting energy expenditure: how to correct mouse metabolic rate for body mass. Nat. Metab., 3:1134–1136, 2021. doi: 10.1038/s42255-021-00451-2.

[18] Plotly Technologies Inc. Collaborative data science. Montreal, QC, 2015. URL https://plot.ly.

