## Supplementary material for "Shiny-Calorie: A context-aware application for indirect calorimetry data analysis and visualization using R": Shiny-Calorie-Supplemental

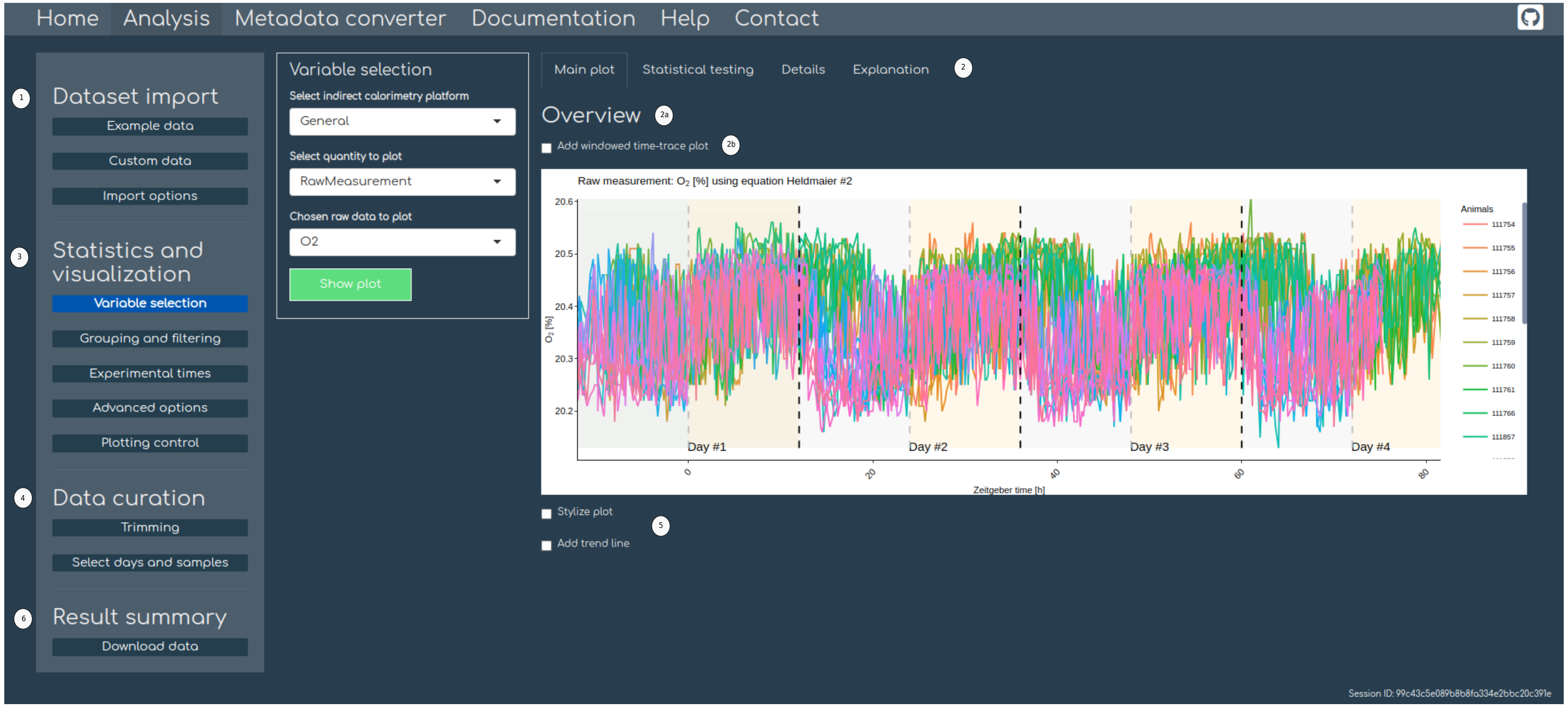


Supplementary Figure S1: **Overview of Shiny-Calorie's user interface.** **1** Dataset import: The section allows to provide own datasets and metadata, or use one of the provided built-in data sets. A user guide is available from the help content in the navigation bar. Under the subsection `Example data` one finds a Ciceron guide which will walk users through the application's features. **2** Shiny-Calorie is subdivided into a left sidebar panel for controls, middle panel for configuration and a main panel on the right for visualization, statistical analysis and explanation of the conducted analyses. **2a** Displays the main view with the selected variable of interest over time and **2b** allows to conduct windowed time analysis, which will be added to the main panel below **5** on the bottom. In the main view there is always the possibility to adjust aesthetics of generated plots by using the `Stylize plot` checkbox. **3** Panel to select a metabolic variable for visualization and analysis, also allows for grouping, filtering, annotation of experimental times and advanced options typically not required in a first analysis. **4** Allows for data curation, e.g. trimming of experimental times and, days and sample selection. Raw data curation is specified during the `Dataset import` in **1** and can be configured via the `Import options` section. **6** Allows to export and download all plots and accumulated data frames of calculated and derived quantities as a single compressed `ZIP` archive. The navigation bar links to documentation and Metadata converter Shiny/R app to assist users to fill out the Metadata Sheet [1].

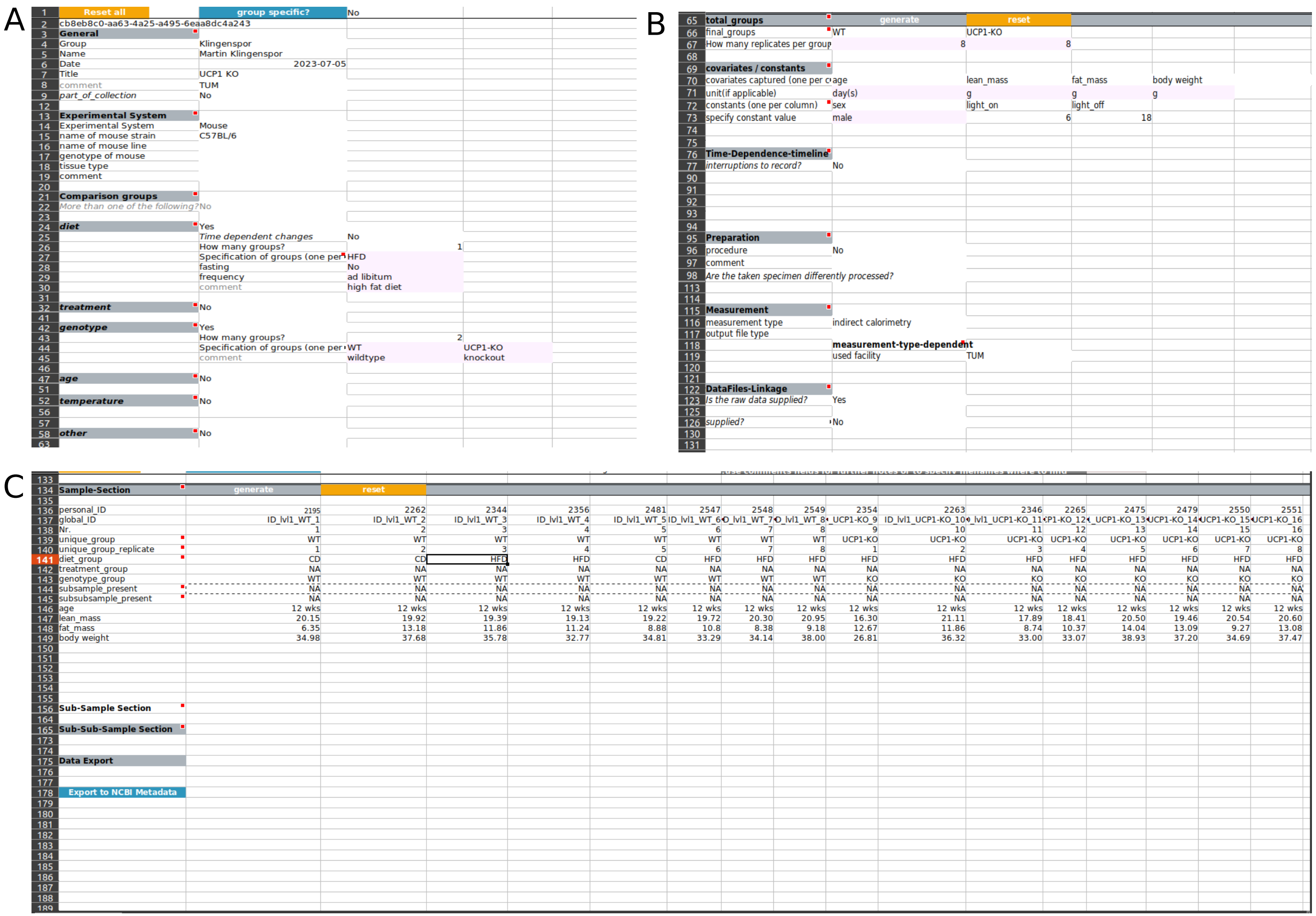


Supplementary Figure S2: **Metadata** as provided through the standardized metadata sheet [1] using a controlled vocabulary provided by the accompanying metadata ontology (not shown). A. Basic description and information of the experiment and experimental setup is given, e.g. diet, treatments, genotype. Other categorical groups can be added *ad libitum* if applicable. B. Specification of measured covariates and constants. C. Compiled sample table for experiment with the possibility for sub-sampling.

| | A | B | C | D | E | F | G | H | I | J | K | L | M |
|---|---|---|---|---|---|---|---|---|---|---|---|---|---|
| 1 | 20200514_SD_Ucpdd_K2 | | TX001 | TX002 | TX003 | | | | | | | | |
| 2 | | TSE LabMaster V6.3.3 (2017-3514) | | | | | | | | | | | |
| 3 | Box | Animal No. | Weight [g] | Diet | Genotype | Text3 | | | | | | | |
| 4 | 4 | 2344 | 35.78 | HFD | WT | | | | | | | | |
| 5 | 5 | 2346 | 31.03 | HFD | KO | | | | | | | | |
| 6 | 7 | 2354 | 33.07 | HFD | KO | | | | | | | | |
| 7 | 8 | 2356 | 32.77 | HFD | WT | | | | | | | | |
| 8 | | | | | | | | | | | | | |
| 9 | Date | Time | Animal No. | Box | S.Flow | Ref.O2 | Temp | O2 | dO2 | dCO2 | VO2(3) | VCO2(3) | RER |
| 10 | | | | | [l/min] | [%] | [°C] | [%] | [%] | [%] | [ml/h] | [ml/h] | |
| 11 | 14.05.2020 | 16:30 | 2344 | 4 | 0,4 | 20,75 | 25,8 | 20,32 | 0,4297 | 0,3708 | 135 | 112 | 833 |
| 12 | 14.05.2020 | 16:35 | 2344 | 4 | 0,4 | 20,74 | 26,8 | 20,39 | 0,3487 | 0,3011 | 109 | 91 | 834 |
| 13 | 14.05.2020 | 16:40 | 2344 | 4 | 0,4 | 20,74 | 27,5 | 20,43 | 0,3094 | 0,2608 | 98 | 79 | 0,81 |
| 14 | 14.05.2020 | 16:45 | 2344 | 4 | 0,4 | 20,75 | 28 | 20,44 | 0,3082 | 0,2618 | 96 | 78 | 817 |
| 15 | 14.05.2020 | 16:50 | 2344 | 4 | 0,4 | 20,75 | 28,4 | 20,45 | 0,2947 | 0,2539 | 92 | 76 | 832 |
| 16 | 14.05.2020 | 16:55 | 2344 | 4 | 0,4 | 20,75 | 28,7 | 20,45 | 0,3057 | 0,2569 | 96 | 77 | 806 |
| 17 | 14.05.2020 | 17:00 | 2344 | 4 | 0,4 | 20,75 | 29 | 20,46 | 0,2885 | 0,2397 | 90 | 72 | 795 |
| 18 | 14.05.2020 | 17:05 | 2344 | 4 | 0,4 | 20,75 | 29,2 | 20,47 | 0,2787 | 0,2298 | 86 | 68 | 788 |
| 19 | 14.05.2020 | 17:10 | 2344 | 4 | 0,4 | 20,75 | 29,2 | 20,5 | 0,2505 | 0,2112 | 78 | 63 | 0,81 |
| 20 | 14.05.2020 | 17:15 | 2344 | 4 | 0,4 | 20,76 | 29,1 | 20,52 | 0,2394 | 0,2028 | 75 | 61 | 815 |
| 21 | 14.05.2020 | 17:20 | 2344 | 4 | 0,4 | 20,75 | 29,1 | 20,48 | 0,2775 | 0,2446 | 86 | 74 | 855 |
| 22 | 14.05.2020 | 17:25 | 2344 | 4 | 0,4 | 20,76 | 29,2 | 20,44 | 0,3192 | 0,2711 | 103 | 84 | 817 |
| 23 | 14.05.2020 | 17:30 | 2344 | 4 | 0,4 | 20,75 | 29,2 | 20,56 | 0,1964 | 0,1596 | 62 | 48 | 774 |
| 24 | 14.05.2020 | 17:35 | 2344 | 4 | 0,4 | 20,76 | 29,2 | 20,57 | 0,1866 | 0,1532 | 58 | 46 | 784 |
| 25 | 14.05.2020 | 17:40 | 2344 | 4 | 0,4 | 20,76 | 29,3 | 20,59 | 0,1682 | 0,14 | 53 | 42 | 797 |
| 26 | 14.05.2020 | 17:45 | 2344 | 4 | 0,4 | 20,76 | 29,3 | 20,59 | 0,1694 | 0,1439 | 53 | 43 | 817 |
| 27 | 14.05.2020 | 17:50 | 2344 | 4 | 0,4 | 20,76 | 29,4 | 20,58 | 0,1805 | 0,1586 | 56 | 48 | 852 |
| 28 | 14.05.2020 | 17:55 | 2344 | 4 | 0,4 | 20,76 | 29,5 | 20,44 | 0,3204 | 0,2853 | 99 | 86 | 866 |
| 29 | 14.05.2020 | 18:00 | 2344 | 4 | 0,4 | 20,76 | 29,7 | 20,49 | 0,2689 | 0,2362 | 84 | 71 | 851 |
| 30 | 14.05.2020 | 18:05 | 2344 | 4 | 0,4 | 20,76 | 29,7 | 20,49 | 0,2738 | 0,2392 | 84 | 71 | 846 |
| 31 | 14.05.2020 | 18:10 | 2344 | 4 | 0,4 | 20,76 | 29,8 | 20,43 | 0,3364 | 0,2942 | 104 | 88 | 847 |

Supplementary Figure S3: **Raw data** export as generated by users from a TSE LabMaster v7/v8 system as `CSV` file (viewed in LibreOffice). Metadata is encoded in header rows (1-8) and separated from raw data by a single empty line. Note that raw data is recorded in a row-oriented format for individual samples (e.g. animals) here, raw data can also be exported into a column-oriented format (not shown).

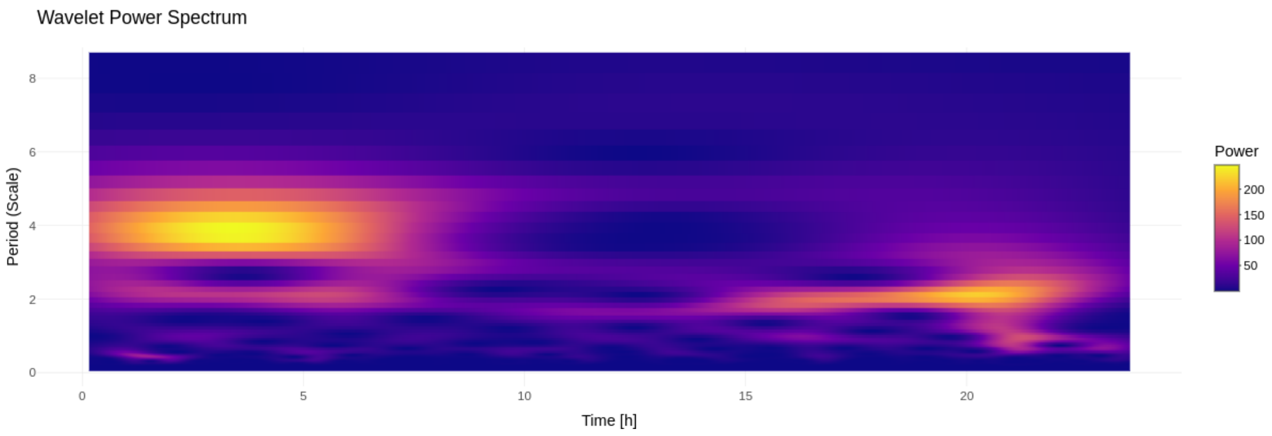


Supplementary Figure S4: **Wavelet power spectrum.** A detailed visualization of the dominant frequency components with changes over time is shown for the signal of metabolic rate of a single individual sample.

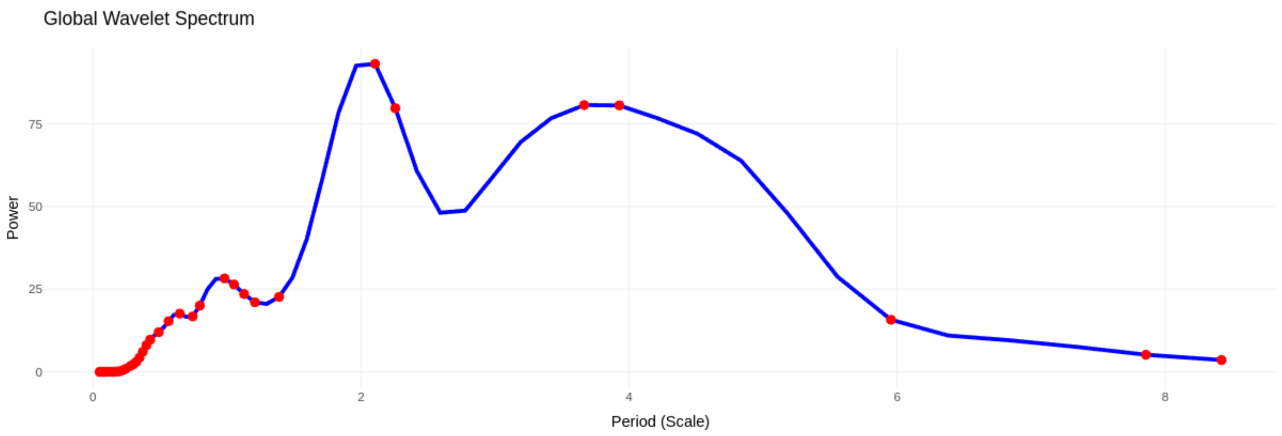


Supplementary Figure S5: **Significant periods in the global wavelet spectrum.** Significant frequency components are visualized which are present in the analyzed non-stationary signal of metabolic rate for a single individual sample.

Supplementary Table S1: **Supported input file formats**. Tabulated are all currently, i.e. in Shiny-Calorie v0.4.7 version, supported file formats as output from the associated metabolic phenotyping platforms. File types are allowed to be combined during upload (multiple cohort study) and their metadata labels are harmonized.

| **Manufacturer** | **File type** | **Supported version** | **Format specification** |
|---|---|---|---|
| Sable Systems | .xlsx | $\geq$ v22.0.0 | Promethion Live |
| TSE Systems | .csv / .tsv | $\geq$ v5.0.0, $\geq$ v6.0.0 | LabMaster |
| TSE Systems | .csv / .tsv | $\geq$ v7.0.0, $\geq$ v8.0.0 | PhenoMaster |
| TSE Systems | .csv | - | Calobox |
| COSMED Platform | .xlsx | - | - |

Supplementary Table S2: **Available heat production equations**. Tabulated are the available heat production equations with definition, units and literature reference. For a discussion of these equations, i.e. how the equations have been derived from experimental data sets and their application and implications, cf. [2].

| **Name** | **Equation** | **Unit** | **Reference** |
|---|---|---|---|
| Heldmaier's first | $(4.44 + 1.43 \times RER) + \dot{V}O_2$ | $\frac{ml}{h}$ | [3] |
| Heldmaier's second | $\dot{V}O_2 \times (6 + RER + 15.3) \times 0.278$ | $\frac{ml}{h}$ | [3] |
| Weir | $16.3 \times \dot{V}O_2 + 4.57 \times RER$ | $\frac{ml}{h}$ | [4] |
| Ferrannini | $16.37117 \times \dot{V}O_2 + 4.6057 \times RER$ | $\frac{ml}{h}$ | [5] |
| Lusk | $15.79 \times \dot{V}O_2 + 5.09 \times RER$ | $\frac{ml}{h}$ | [6] |
| Elia | $15.8 \times \dot{V}O_2 + 5.18 \times RER$ | $\frac{ml}{h}$ | [7] |
| Brouwer | $16.07 \times \dot{V}O_2 + 4.69 \times RER$ | $\frac{ml}{h}$ | [8] |

### Supplementary material for Shiny-Calorie: A context-aware application for indirect calorimetry data analysis and visualization using R

Stephan Grein[1] Tabea Elschner[2,3,†] Ronja Kardinal[4,†] Johanna Bruder[5] Akim Stromeyer[5] Karthikeyan Gunasekaran[6] Jennifer Witt[6] Hildigunnur Hermannsdottir[7] Janina Behrens[6] Muuez U-Din[8] Jiangyan Yu[9] Gerhard Heldmaier[10] Renate Schreiber[11] Jan Rozman[12] Markus Heine[6] Ludger Scheja[6] Anna Worthmann[6] Jörg Heeren[6] Dagmar Wachten[4] Kerstin Wilhelm-Jüngling[2,3] Alexander Pfeifer[13] Jan Hasenauer[1,*] Martin Klingenspor[5,7,14,*]

**1 Life and Medical Sciences (LIMES) Institute and Bonn Center for Mathematical Life Sciences, University of Bonn, Bonn, Germany**
**2 Institute for Cardiovascular Sciences, University Hospital, University of Bonn, Bonn, Germany**
**3 Institute for Neurovascular Cell Biology, University Hospital, University of Bonn, Bonn, Germany**
**4 Institute of Innate Immunity, University Hospital, University of Bonn, Bonn, Germany**
**5 EKFZ - Else Kröner-Fresenius Center for Nutritional Medicine, Technical University of Munich, Freising-Weihenstephan, Germany**
**6 Department of Biochemistry and Molecular Cell Biology, University Medical Center Hamburg-Eppendorf, Hamburg, Germany**
**7 Chair of Molecular Nutritional Medicine, TUM School of Life Sciences, Technical University of Munich, Freising, Germany**
**8 Turku PET Centre, University of Turku and Turku University Hospital, Turku, Finland**
**9 Institute of Clinical Genetics and Genomic Medicine, University Hospital, Würzburg, Germany**
**10 Animal Physiology, Department of Biology, Marburg University, Marburg, Germany**
**11 Institute of Molecular Biosciences, University of Graz, Graz, Austria**
**12 Luxembourg Centre for Systems Biomedicine, University of Luxembourg, Esch-Belval, Luxembourg**
**13 Institute of Pharmacology and Toxicology, University Hospital, University of Bonn, Bonn, Germany**
**14 ZIEL - Institute for Food & Health, Technical University of Munich, Freising, Germany**

### 1 Introducing Shiny-Calorie

Shiny-Calorie, a context-ware application for indirect calorimetry data analysis and visualization using R, is an open-source, reactive web application for the comprehensive visualization and statistical analysis of general indirect calorimetric data sets arising from indirect calorimetry experiments using common metabolic phenotyping platforms, i.e. TSE Systems Phenomaster/LabMaster, Sable Systems Promethion and the COSMED platform, cf. Tab. S1. The previously referenced table contains a list of supported file formats and versions for the mentioned metabolic phenotyping platforms. Shiny-Calorie ingests data and standardized metadata, and automatically integrates and harmonizes the data sets for analysis. For an overview of Shiny-Calorie's user interface, cf. Fig. S1.

**Usage of the web application** The Shiny-Calorie web application is deployed on two redundant resources, on our own infrastructure `https://shiny.iaas.uni-bonn.de/Shiny-Calorie` and on the infrastructure as service (IaaS) platform ShinyApps.io `https://shinyapps.io/stephanmg/calorimetry`.

Common web browsers in recent versions are suggested (e.g. Chrome/Chromium, Firefox), since Shiny/R inherently makes use of the web technology stack, e.g. CSS, JS, etc., for rendering of content and user interaction.

Users interface through the browser to load, analyze data and generate visualizations of indirect calorimetry experiments and conduct statistical hypothesis testing. Two published ready-to-use example datasets (UCP1-KO and DAKO study) are integrated into the web application (datasets are loadable via two different buttons indicating the dataset directly on the application's landing page) for a quick exploration of the functionality of Shiny-Calorie. The same datasets are detailed and analyzed in Shiny-Calorie's accompanying documentation on `https://ICB-DCM.github.io/Shiny-Calorie/`. Additionally, users can compile their custom datasets by use of our calorimetry tools (using the Apache/Solr2 REST API) accessing the International Mouse Phenotyping (IMPC) database, see Section 6.3 on IMPC dataset generation.

*Nota bene*: To support efficiently multiple users in their analysis requirements, we utilize a custom lab-internal small-scale Kubernetes cluster (4 worker nodes and 1 master controller node). This allows us to load balance and upscale on demand to support efficient usage of the web application and adapt to growing number of users if necessary.

### 2 On-premise installation and deployment

The Shiny-Calorie web application is made available through a containerized OCI docker image via `hub.docker.com`, cf. repository `https://hub.docker.com/r/stephanmg/Shiny-Calorie` for details. Docker images can be accessed via the common workflow, i.e. `docker pull stephanmg/Shiny-Calorie` to obtain the latest tagged version.

Typically Shiny-Calorie will be placed behind a reverse-proxy solution for load balancing with e.g. Nginx on local compute infrastructure (to allow for efficient parallel usage by many different users) or in a setup involving Kubernetes. Start scripts are provided in the source code repository on Github, and an example configuration to set up load balancing via Nginx for multiple parallel running app instances to support efficient multi-user usage of the web application is provided. Thus, Shiny-Calorie can be installed easily on either local computers as well as on cloud infrastructures respectively IaaS providers by means of Docker or drop-in replacement as Podman or the analogues Singularity/Apptainer.

### 3 Installation on a personal computer

For Shiny-Calorie builds are created nightly, i.e. installers for Windows, OSX and Linux (Ubuntu) are available as artifacts from the Github Actions workflows on Shiny-Calorie's Github repository. Standalone versions are provided for situations where connectivity is limited, i.e. in offline field experiments or shielded lab rooms with low or no connection strength for wireless transmission where also no Ethernet is available.

### 4 Code availability

The code of Shiny-Calorie is available in the corresponding Github repository, and installation from source is possible with dependencies managed through `renv`, rendering a native installation of Shiny-Calorie convenient for developers and promotes reproducibility. The corresponding `renv.lock` file is contained within the repository itself, listing required packages and dependencies.

### 5 Documentation

The application provides built-in documentation, see top right corner in the web application. A walk-through, respectively, a guide is available at the user's discretion ('Guide' button in web application) on the web application's landing page. Further documentation is available on the RTD documentation of Shiny-Calorie.

### 6 Application examples

In the following sections, two application examples illustrate the common workflows of analysis for indirect calorimetry experiments with example datasets. The examples are intended to guide new users and to become familiar with the features, layout of the user interface and the functionality of Shiny-Calorie. These examples were developed in close collaboration with wet-lab scientists and are based on routinely applied standard operating procedures (SOPs) and analysis protocols in the labs. The methods implemented in Shiny-Calorie automatize repetitive day-to-day workflows by a joint integration of data and metadata as recorded in the lab book using the Metadata sheet.

Additional guidance and documentation are available through the built-in help system and step-by-step tutorial embedded within the application's user interface. Furthermore, two additional use-cases respectively examples are described in tutorial style on Shiny-Calorie's documentation website: `https://ICB-DCM.github.io/Shiny-Calorie`

#### 6.1 Dataset I: Disentangle activity-related energy expenditure from resting metabolic rate and statistical hypothesis testing on data stratifications

To illustrate the workflow of data analysis after an indirect calorimetry experiment has been conducted, we consider the case of reconstructing energy expenditure (EE) and analysis thereof and perform additional subsequent downstream analysis tasks, i.e. calculation of the resting metabolic rate (RMR), total energy expenditure (TEE) and the activity-related energy expenditure (AEE) stratified by genotype. We consider here a stratification with respect to the genotype, which is a specific choice made to illustrate the dataset. It would also be possible to group by diet, condition, temperature and related variables.

Prior to initiating any analysis in Shiny-Calorie, the raw input data—acquired from one of the supported metabolic phenotyping platforms (MPPs)—must successfully pass the built-in consistency checks. Shiny-Calorie enables the analysis of indirect calorimetry datasets from multiple experiments or cohorts, which can be stratified by variables such as genotype or treatment condition (e.g., cold exposure), or analyzed jointly in multi-factorial designs. By default, metadata are extracted from the headers of commonly supported file formats. For enhanced standardization and richer annotation, users are encouraged to supply a structured Metadata Sheet [1].

In the following, we outline the individual steps:

1. **Dataset import:** The user-provided indirect calorimetry datasets are imported into the application using the upload dialog. The supported file types and formats are listed in Tab. S1. Labels of different datasets are harmonized automatically without user intervention. The application outputs a warning if not fully-supported datasets are provided and restricts the users from further analysis unless explicitly ignored by the user.

File upload dialog for data sets:

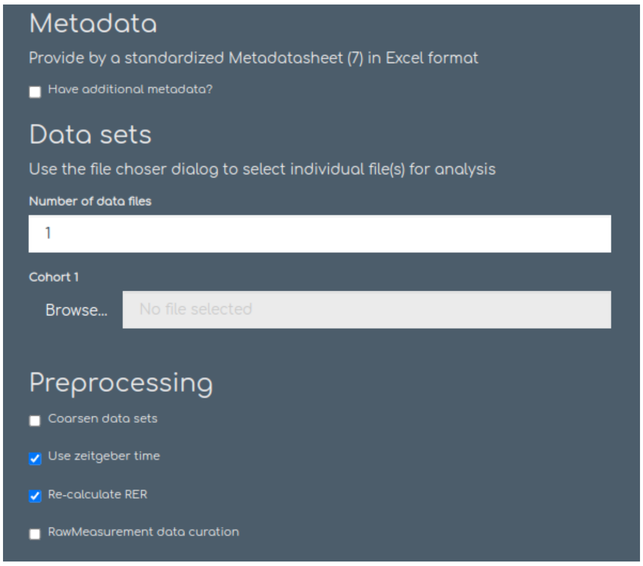


During data import, the user can either flag or remove implausible values through invoking the raw data curation panel. Pre-processing of raw datasets is thus either automatically handled or flagged for manual inspection by users. Pre-processing allows to coarsen the dataset (i.e. short measurement intervals), interpolate to a regular time grid (if unequal measurement interval lengths are given), convert to zeitgeber time (align with the photoperiod) or keep regular calender time. We allow to recalculate the respiratory exchange ratio too, as we observed that some platforms do not correctly output the quantity in `CSV` files occasionally.

Checking the box `RawMeasurement data curation` displays additional panels which various options for outlier removal and marking of outliers, and enables the possibility to activate the raw data consistency checks.

Pre-processing panel:

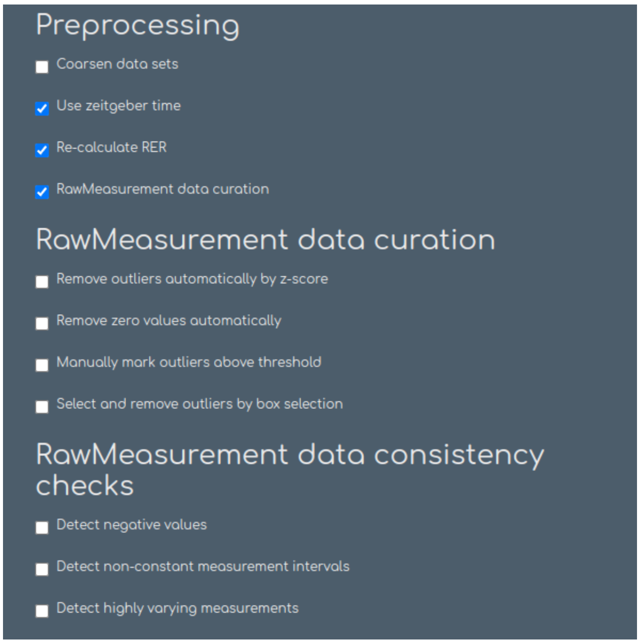


2 **Review of metadata:** The users review the metadata attached to their indirect calorimetry experiments. Purposefully Shiny-Calorie outputs a comprehensive overview of metadata to allow for a detailed analysis of metadata, numerical quantities are output as density plots and categorical data as boxplots. Users compare all metadata and perform multi-way ANOVAs in the `Statistical testing` panel to check consistency of experimental design (and for biases), e.g. even distribution of diets or genotypes, consistent age of samples, or to check for any significant differences in, e.g., body weight which might distort statistical analysis when correcting for body weight to avoid biased analyses.

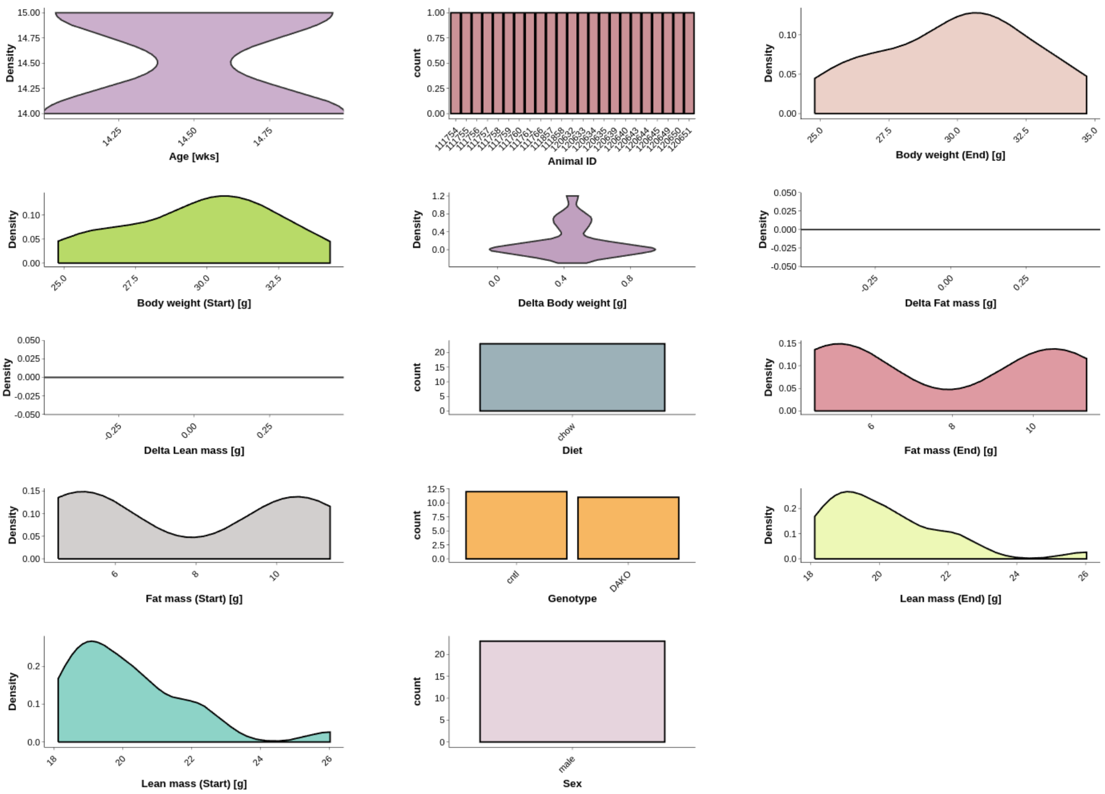

Following initial data visualization and statistical testing, users should ensure that no major biases exist in the experimental design before proceeding with additional quality control steps. This includes the routine inspection of respiratory gas exchange traces ($CO_2$ and $C_2$) and the respiratory exchange ratio (RER), which serve as critical indicators of measurement integrity in indirect calorimetry experiments which is calculated as the fraction of the former quantities.

The metadata review acts as an initial checkpoint to confirm consistency and completeness of the experimental setup. Raw data inspection involves not only assessing the gas exchange profiles and RER, but also, where available, evaluating additional parameters such as temperature and photoperiod information. These supplementary variables help identify inconsistencies or anomalies in the exported data from the metabolic phenotyping platforms.

Statistical testing for categorical metadata:

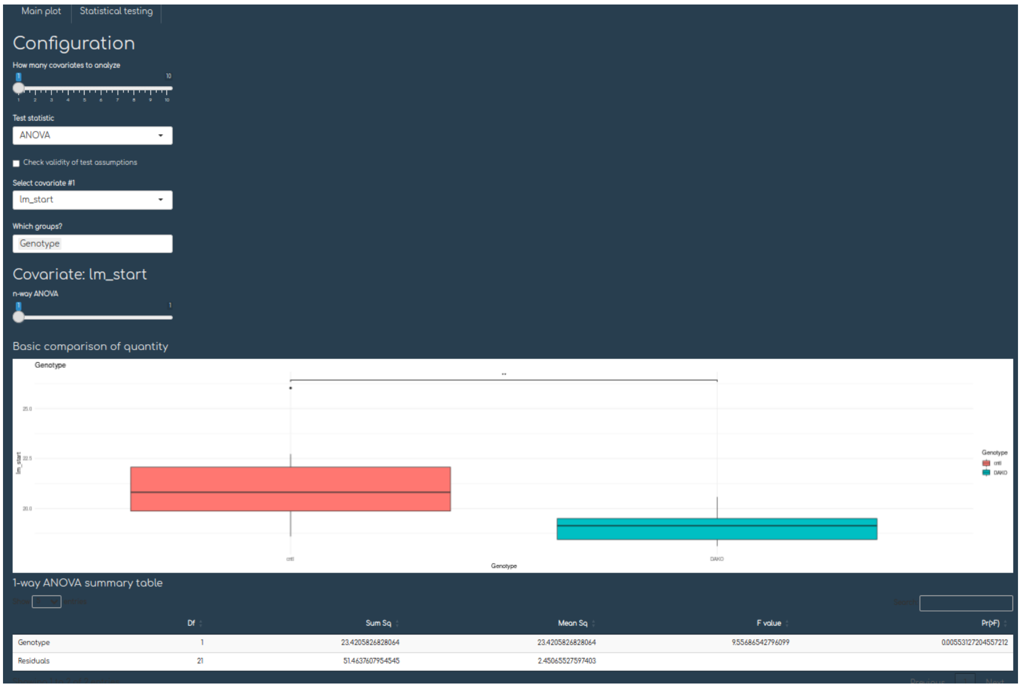


The consistency of input datasets is vital, and thus should always be verified prior to any calculations of EE, analysis, or statistical hypothesis testing thereof. The first important step is concerned with the inspection of the raw oxygen and carbon dioxide signals (saturation and actual production and consumption), as well as derived quantity i.e. the respiratory exchange ratio (RER) mentioned earlier. These quantities should be examined for inconsistencies or irregularities.

The respiratory exchange ratio (dimensionless) derived measurement visualized over time course of experiment can give important information about substrate utilization and soundness of the indirect calorimetry experiment:

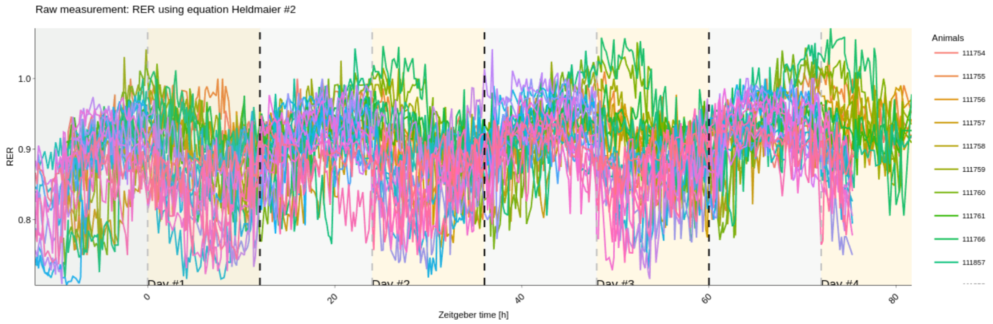

Furthermore, the inspection of the raw oxygen and carbon dioxide signals can help detect experimental flaws or issues with subjects in the calorimetric cage or chamber.

Users can request that subsequent analysis should be prohibited when high-frequency (non-physiologically) measurements or non-plausible input data (e.g. negative values) are detected, resulting in a visual pop-up warning in the web application with additional information in the `Data set loading` panel, cf. Fig. S1 section 1. The warning can be overridden at the discretion of the user.

Plot of oxygen consumption ($VO_2$) as a raw measurement visualized over time course of experiment:

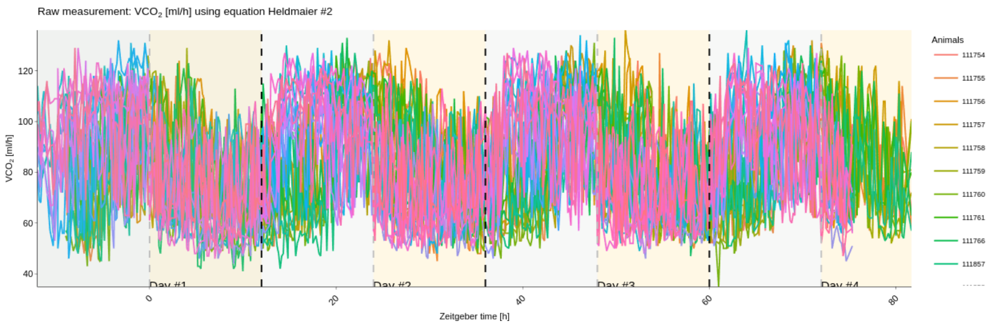


Plot of carbon dioxide production ($VCO_2$) as a raw measurement visualized over time course of experiment.

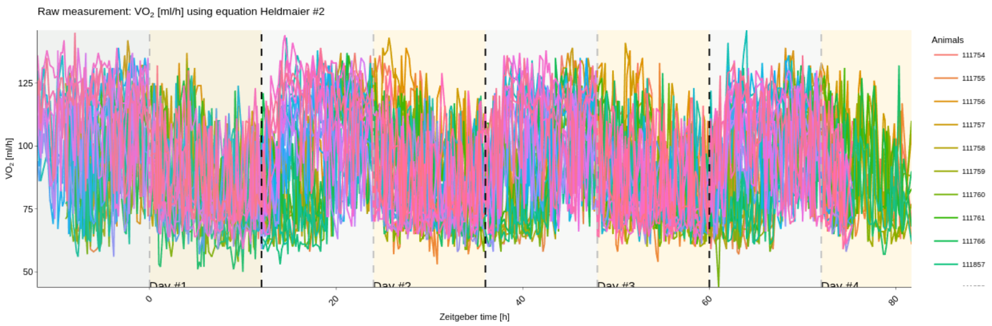


Additionally, the signals of $O_2$ and $CO_2$ saturation should be inspected. A visualization of these quantities can reveal if there are any general issue with the indirect calorimetry experiment, i.e. air flows within the metabolic cages, or an issue with the MPP as set up in the laboratory environment. Empty control cages can be excluded from analysis as well.

Plot of the raw measurement oxygen saturation:

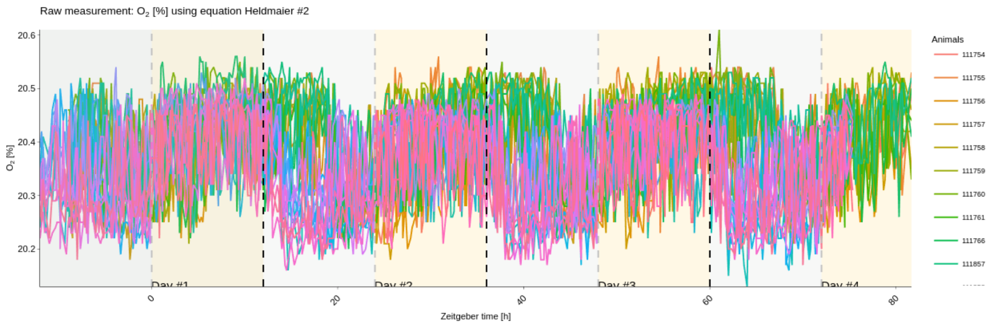


Plot of the raw measurement carbon dioxide saturation:

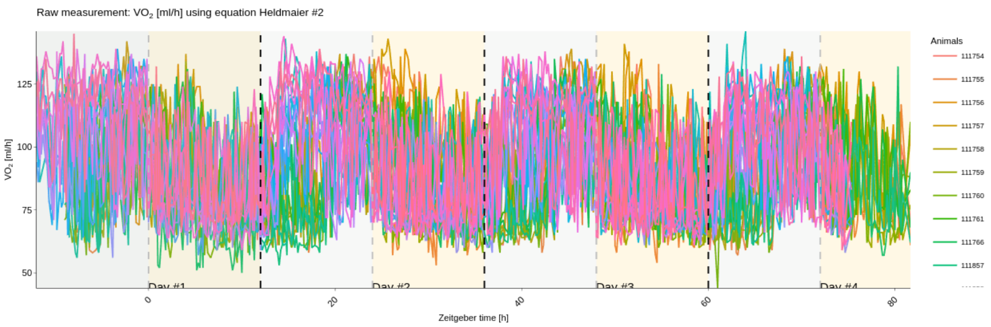


3 **Data curation:** If inconsistencies or irregularities in datasets have been detected, the user can request the removal of these outliers (e.g. exclude sick or erratic animals), trim considered experimental times from the start or end of the indirect calorimetry experiments or select a subset of given experimental day only. Trimming of data might be necessary because of the time-dependent accommodation and acclimatization of animals in metabolic cages. Opening the cage or handling the laboratory animals, distorts the $O_2$ or $CO_2$ signal (or derived quantities i.e. RER) in the early and late stages of an experiment.

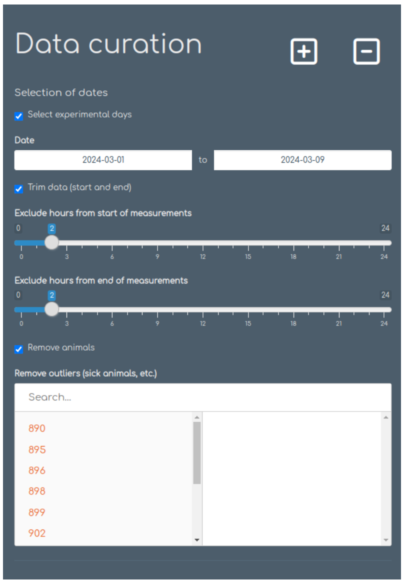


Without correcting for experimental flaws or implausible values, subsequent downstream analysis will receive eventually inconsistent input data and results and statistical analysis are biased. Date ranges are specified via the data input ranges for start and end of experiment. Hours at the beginning or end of an experiment can be trimmed by utilizing the provided sliders to trim from start and end respectively. Sick or erratic animals can be excluded by selecting animal identifiers (sample IDs) in the selection menu. Note data curation is available for all visualizable quantities which users can select in Shiny-Calorie.

4 **Reconstruction of energy expenditure**. The user selects a heat production equation for their experiments. Commonly used heat production formulas to reconstruct energy expenditure and derived quantities are available in Shiny-Calorie (Tab. S2). By a modular design additional heat production formulas can easily be integrated by R users. For exploratory testing, the user can chose two heat production formulas and test for differences in heat production based on the choice they made in an energy expenditure heat production formulas scatter plot. The chosen heat production formula will be used during all downstream analysis and visualization tasks and can be overridden with this single configuration option to allow for a consistent and transparent analysis. In general quantities can be grouped or filtered by metadata, not only for energy expenditure but other quantities.

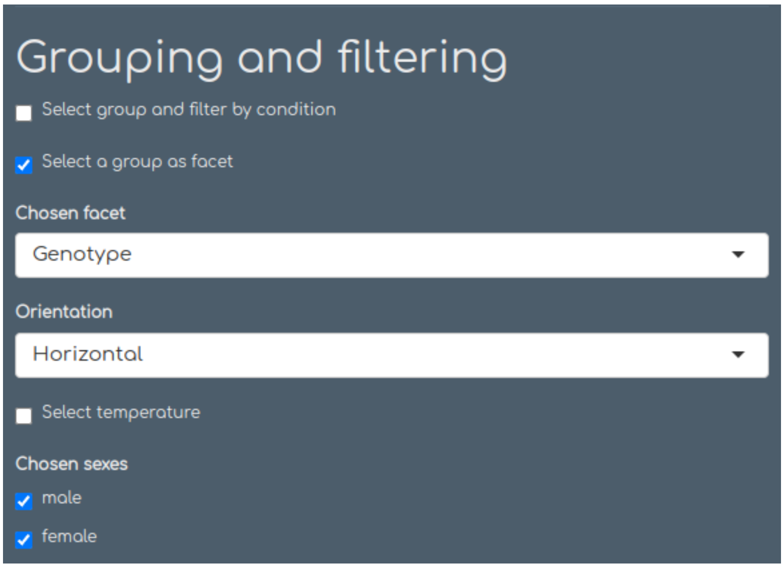


**Grouping and filtering**. Any raw, calculated or derived quantity can either be filtered by using a level or grouped by facets (factors) based on imported metadata. Further samples can be filtered for temperature and based on sex.

*Nota bene*: The choice of a specific heat production formulas had empirically neither a pronounced nor significant difference on energy expenditure as calculated from our test datasets using the raw curated or non-curated measurements from the indirect calorimetry experiment. The user can determine the effects of different heat production equations for energy expenditure calculation using their dataset by using the plotting routine `CompareHeatProductionFormulas` from the `Variable selection` section by selecting the`General` indirect calorimetry platform.

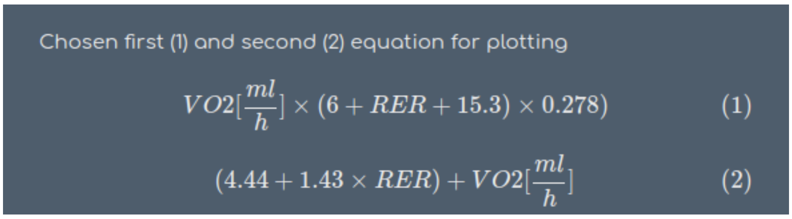


**Heat production equation**. Choosing from the available heat production equations for energy expenditure calculation from respiratory gas exchange data, see also Tab. S2 for options.

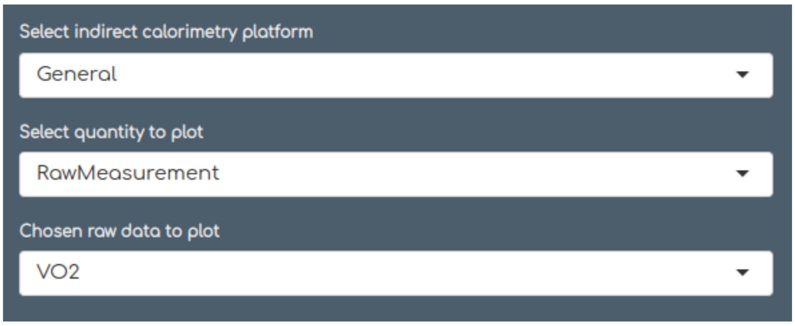


**Selection of analysis**. Choose a raw quantity, derived quantity or energy expenditure quantities to visualize.

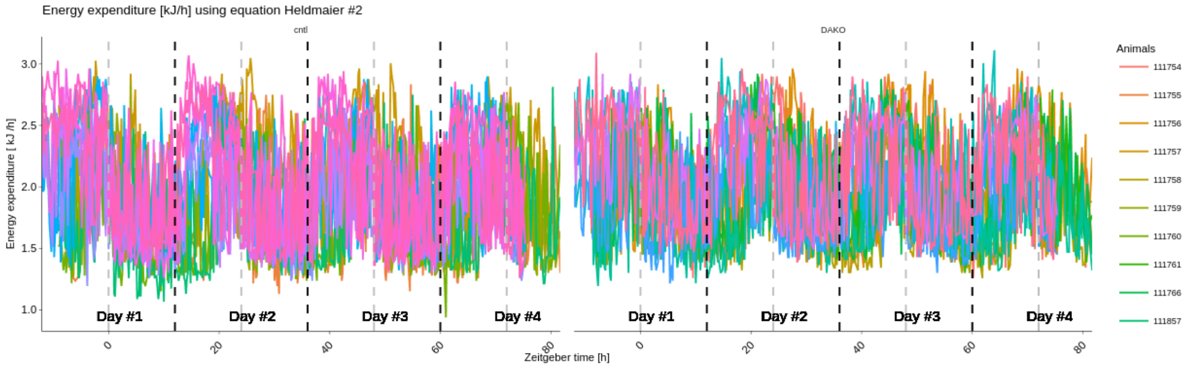


**Energy expenditure over time stratified by the condition diet**. Individual energy expenditure traces of animals are shown and can interactive be controlled in plots with the Plotly library.

5 **Check point.** After all of the previous steps do not detect irregularity or inconsistencies, neither visually nor through statistically testing, users can proceed with downstream analysis.

6 **Calculation of derived quantities**. Resting metabolic rate, total energy expenditure and activity-dependent energy expenditure can be visualized as well.

RMR stratified by genotype for each individual subject:

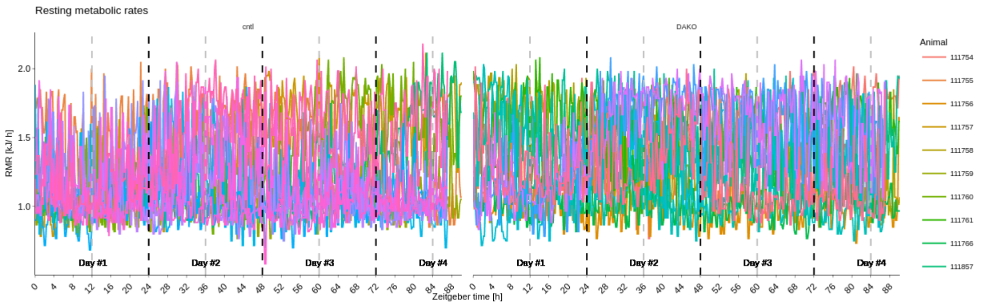


TEE stratified by genotype for each individual subject:

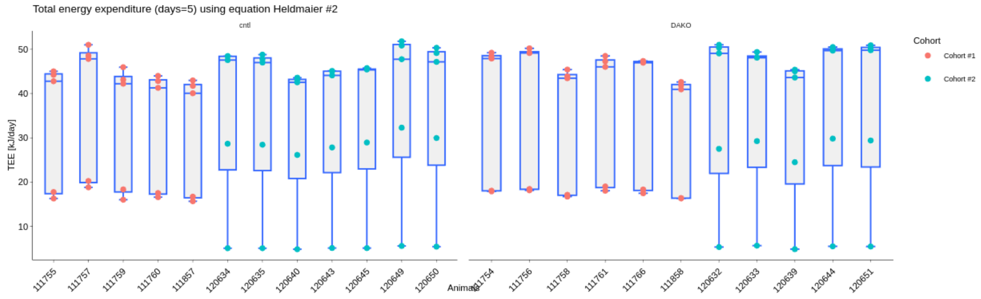

Separation of AEE and RMR:

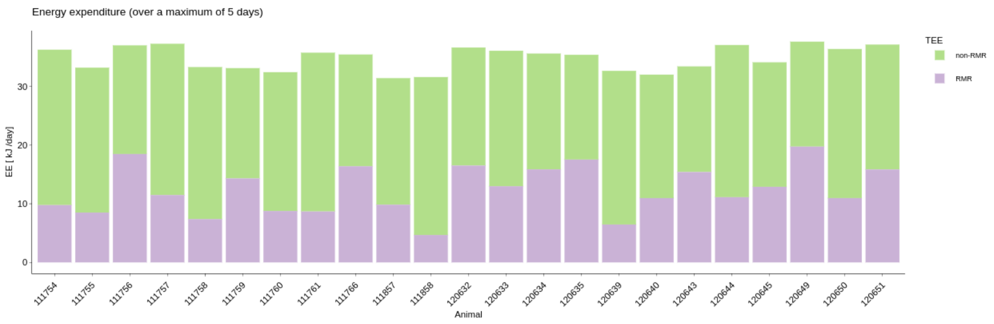


Shiny-Calorie allows also for trend detection, by estimation of the mean traces for all metabolic variables (using GAMs). Mean traces for RMR stratified by genotype:

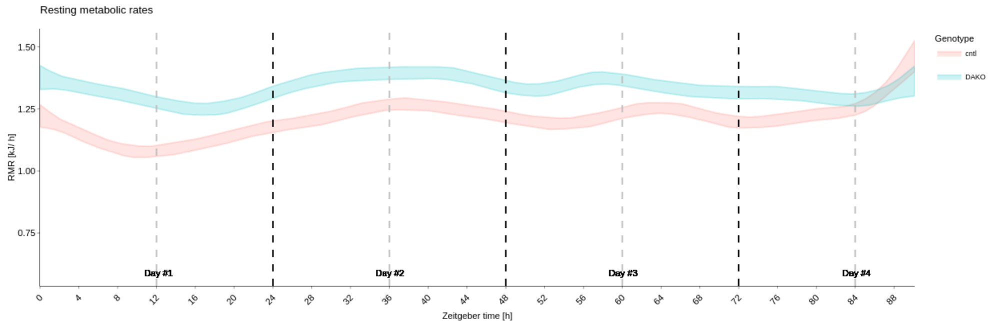


7 **Statistical analysis:** A typical analysis case in indirect calorimetry experiments is concerned with a discrimination of genotype or diet effect on RMR and TEE (and which amount is related to AEE), i.e. energy expenditure stratified across conditions (genotype, diet or both) is of interest to the researchers. In this application example, we compare genotypes (wildtype (WT) and knockout (KO)) with identical diets for an all-male cohorts experiments recorded with the TSE metabolic phenotyping platform.

Thus, the researcher is left with the following steps:

- Calculate total energy expenditure: TEE
- Calculate resting metabolic rate: RMR
- Contrasting and statistical testing of TEE, AEE and RMR

After calculation of TEE and RMR (also termed non-activity related energy expenditure) the following hypothesis should be answered: Is there a genotype effect in RMR or TEE depending on the WT or KO genotype for these cohorts. To answer these questions, the user can conduct a 1-way ANCOVA or 2-way ANCOVA. When a second independent grouping variable, e.g. diet information, becomes or is available through the provided metadata sheet, multi-factorial designs are available to the users. For a visualization of TEE (or EE) and RMR quantities as boxplot or scatte rplots with regression lines, see below.

Boxplot comparison of TEE stratified by genotype, p-values are reported based on ANOVA with Wilcoxon-test.

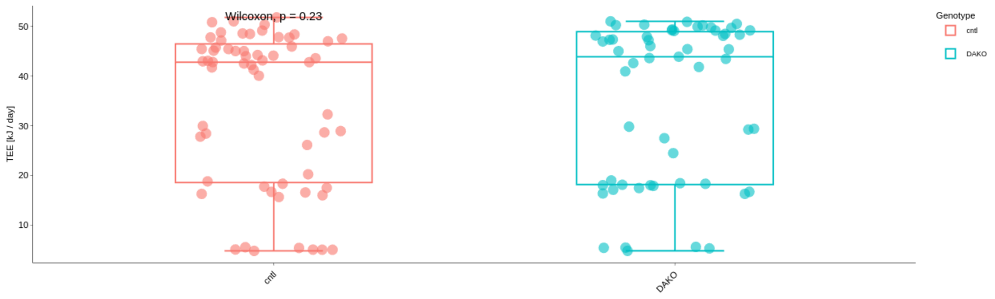


A scatter plot with regression lines for each genotype, $R^2$-values are reported:

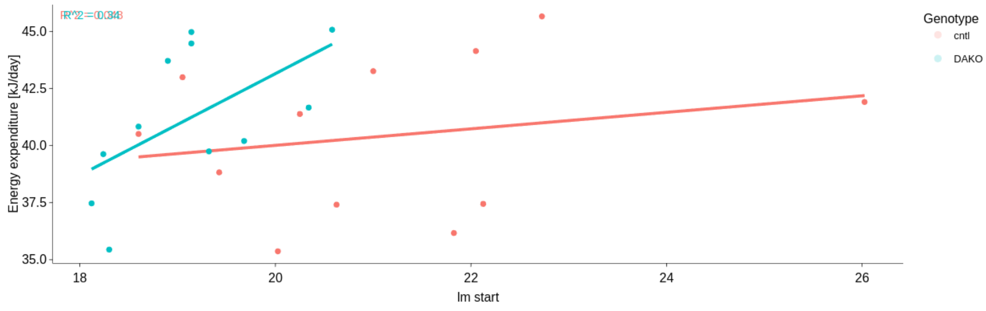


Prior to outputting statistics and conducting of statistical testing, test assumptions, e.g. normality of residuals homoscedasticity of data are verified through visuals, e.g. diagnostic plots, in the application and p-values are reported following. The user can switch between parametric and non-parametric testing depending on the properties of the data set.

When test assumptions and significance levels are met as required by the corresponding tests, conclusions can be drawn and data sets and high-quality figures exported.

1-way ANCOVA comparing KO and WT genotype corrected for body weight variation reveals no difference in RMR and in agreement with previously published results [9].

Test assumption for parametric testing (ANOVA/ANCOVA), e.g. normality of residuals, homoscedasticity and homogeneity of regression slopes are checked and reported to the user.

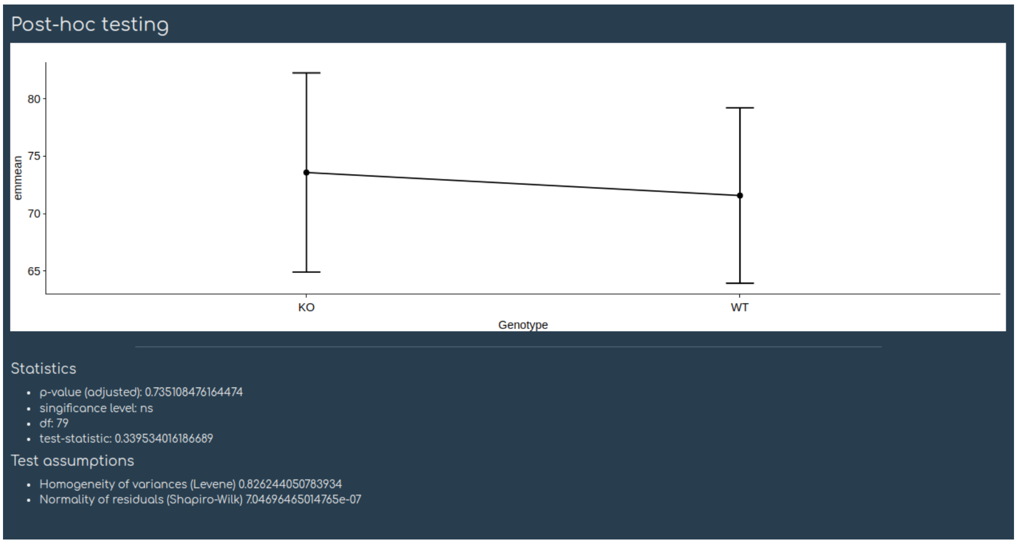


To check for applicability (validate test assumptions for parametric testing) of a statistical test, the corresponding checkbox should be marked:

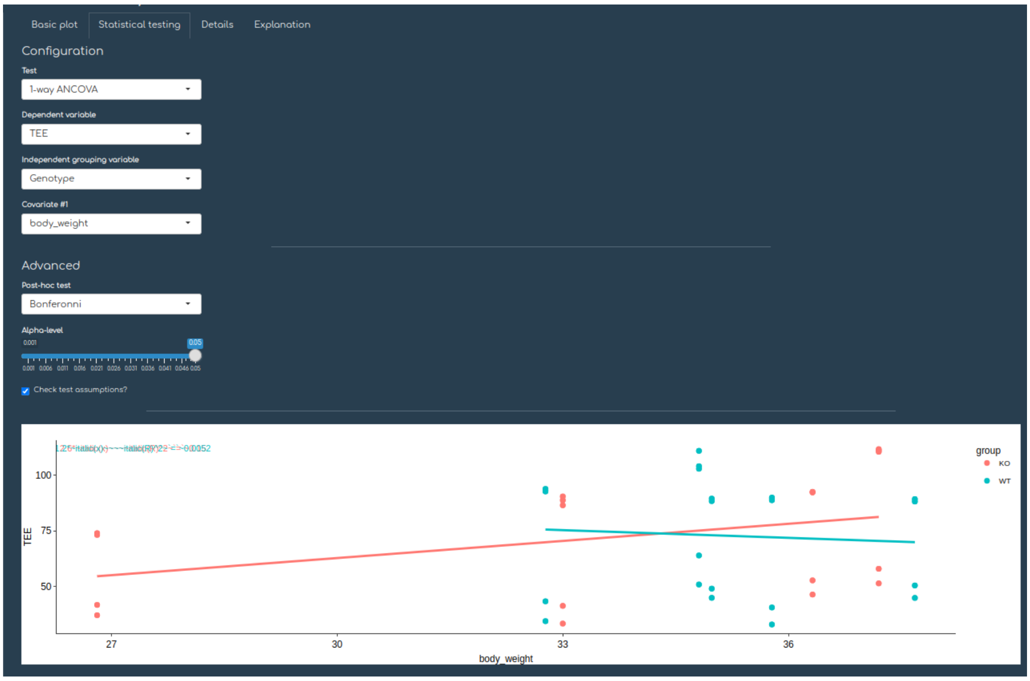


Estimated marginal means for an exemplary 2-way ANCOVA:

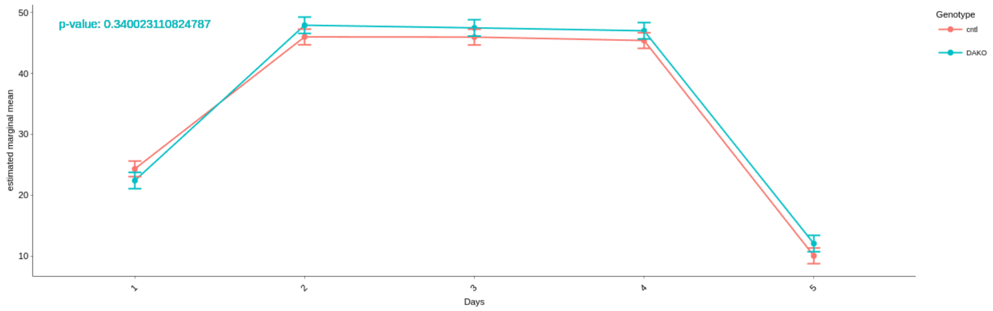

Box plots for the 2-way ANCOVA with additional grouping by Days:

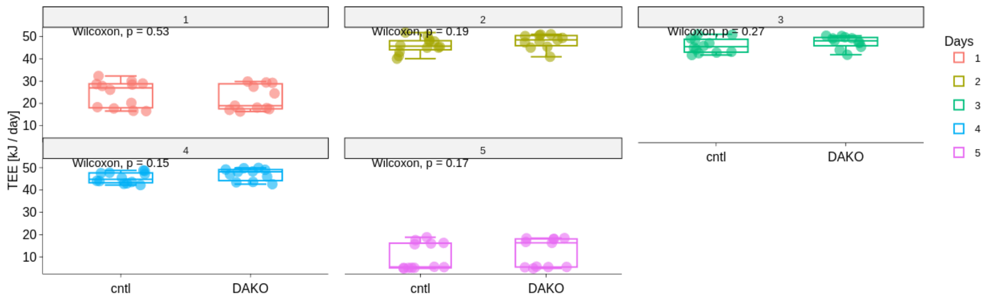


Typically ANCOVA is performed with whole body weight as a covariate we require to correct for, however, by the availability of additional metadata, it is useful to correct for either lean, fat or the combination of lean and fat body mass instead, as is done recently in increasing numbers for metabolic phenotyping experiments. Furthermore TEE and RMR budgets can be visualized, concluding the analysis of TEE and RMR.

In addition windowed time-trace plots can be generated which display per window (e.g. 5, 10, 20 or 30 minutes) averaged quantities of samples. The windowed time-trace plots render useful, as otherwise grand averages might hide significant differences in the course of a day. In the following plot, statistically significant differences are denoted by an asterisk (*).

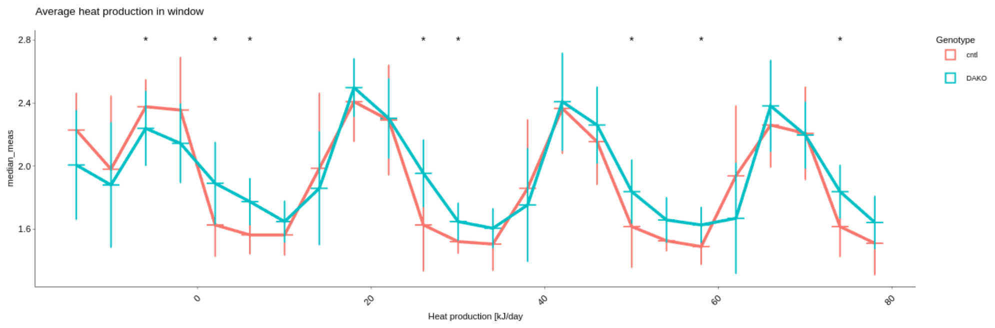


Windowed-time trace analysis reveals, that dependent on the time of day genotype differences can be detected, contrary to analyzing only day-averaged quantities.

#### 6.2 Dataset II. Investigation of locomotion and locomotion budget as a proxy for soundness of experimental setup

It is imperative, whenever locomotion data is available, to verify the behavior of animals in metabolic cages to identify irregularities, idiosyncrasies or problems in experimental setup or with animals, or problems arising during the course of the metabolic phenotyping experiment, which could bias analysis of energy expenditure.

Some metabolic phenotyping platforms, e.g. Sable systems (and also recent versions of TSE Systems. i.e. PhenoMaster), record a subset of locomotion data (steps and time spend in a certain location in the cage by recording time and position (in the (Y, X)-plane using cartesian coordinates) or by recording utilization of the spinning wheel installed in the metabolic cage. A valuable tool therefore is to analyze plots of locomotion density maps (probability density maps), which then can be used to inspect respectively screen for potential issues in indirect calorimetry experiments. Also, the locomotion can be attributed to certain predefined types of physical activity, e.g. drinking, food or using the spinning wheel. A behavioral analysis can be conducted and the locomotion budget can be tabulate and visualized through stacked bar plots. Gained insight can be used for downstream analysis (or in data curation to remove outliers) and the consistent quantification of energy expenditure into the metabolic components activity-related energy expenditure and resting metabolic rate.

Percentage of different locomotional activities (with labels defined as in the Sable/Promethion system) stratified by animal IDs:

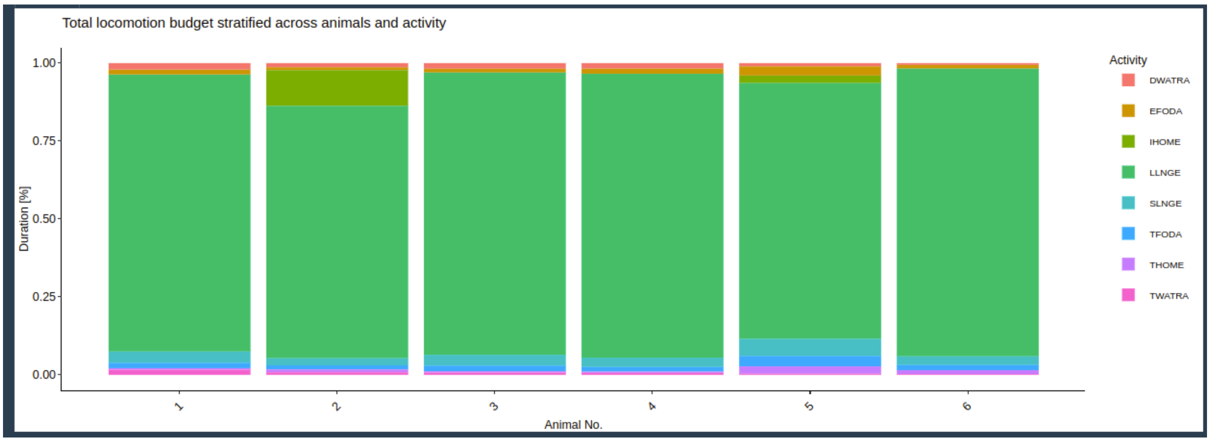


Additionally, not only the locomotor activity can be analyzed but also the budget of locomotion be summarized. In particular a probability density map of animal position in the metabolic cage can be visualized for the purpose of experimental validation. Animal positions are calculated from indirect calorimetry data measuring position of animals at time $t$ in the $(Y, X)$ plane. Dimensions of food hopper (as specified in metadata by Sable/Promethion in Excel workbook), running wheel and water bottle are indicated by colored rectangles.

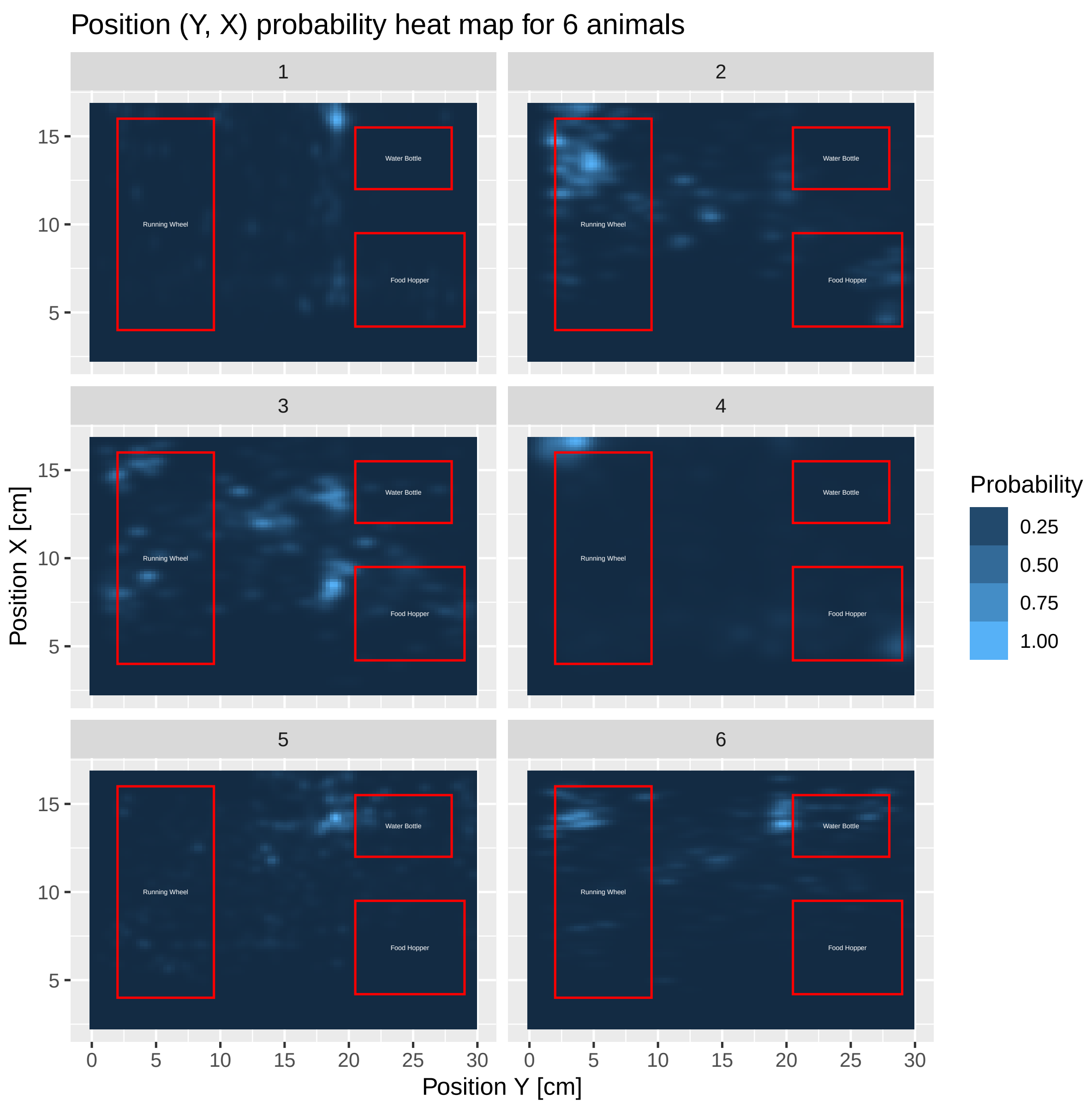


To explore in more depth the dataset and behavior of animals, perhaps to identify issues in experimental setup as mentioned in the previous paragraph, Shiny-Calorie allows for the comprehensive visualization and analysis of related quantities e.g., feeding, drinking, individual step counts, odometer or tracking of body weight. Body weight tracking shows stepwise (discrete) increases and decreases, because animals only sporadically use the scale in the metabolic cage to allow for the measurement of the total body weight.

Step counts are discrete and determined by animals walking through laser light barriers, similarly the distance traveled is determine by taking into account the time between two events of breaking the barrier.

Additionally some setups use a spinning wheel to allow for physical activity of animals, which is measured as another covariate (discrete step counts). Drink and food can be monitored quasi continuously from the provided food in the hamper and water in the drinking bottle. Similarly feeding and drinking periods can be summarized to use the information stored in the IC dataset comprehensively.

Cumulative discrete count of steps in $(Y, X)$ plane:

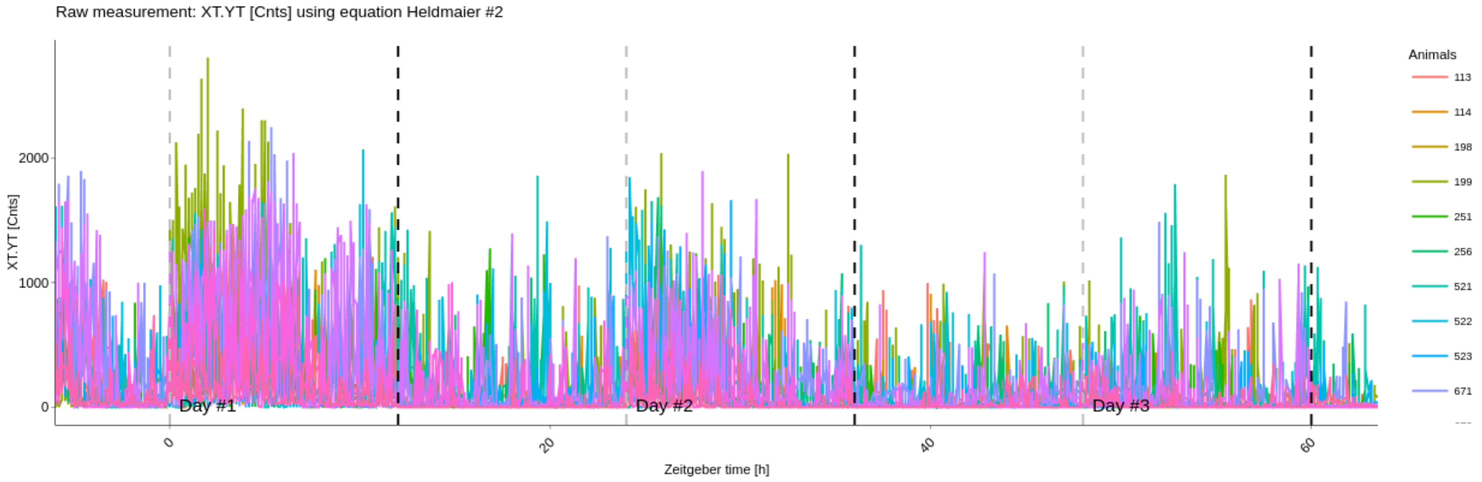


Odometering. Cumulative distance traveled in a metabolic cage per animal:

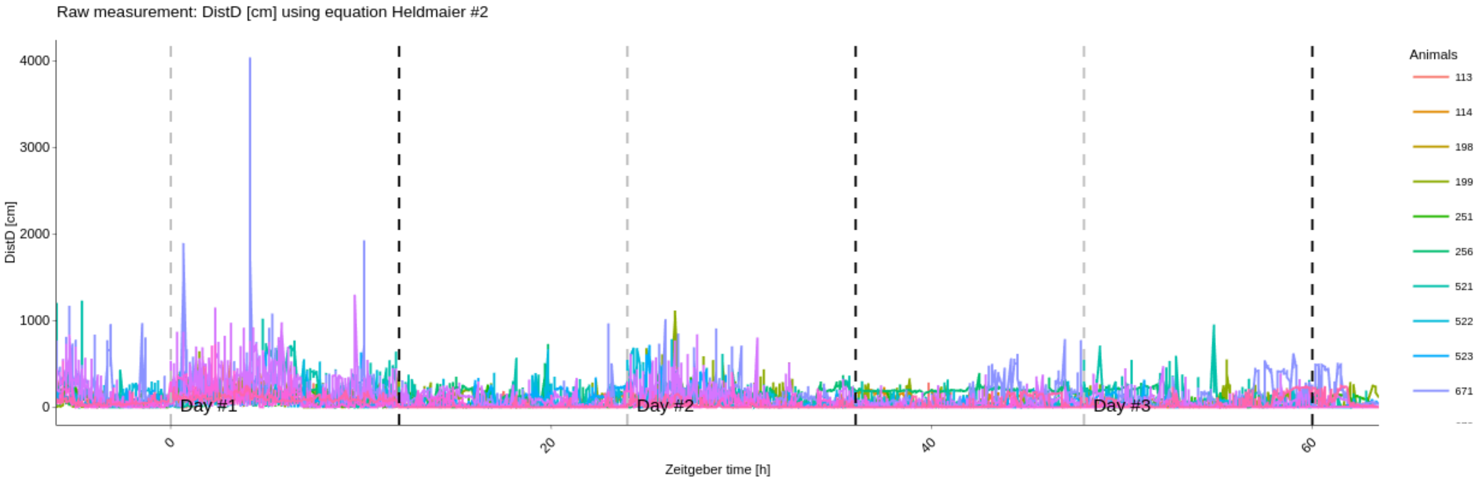

Feeding periods. Cumulative amount of food intake per animal:

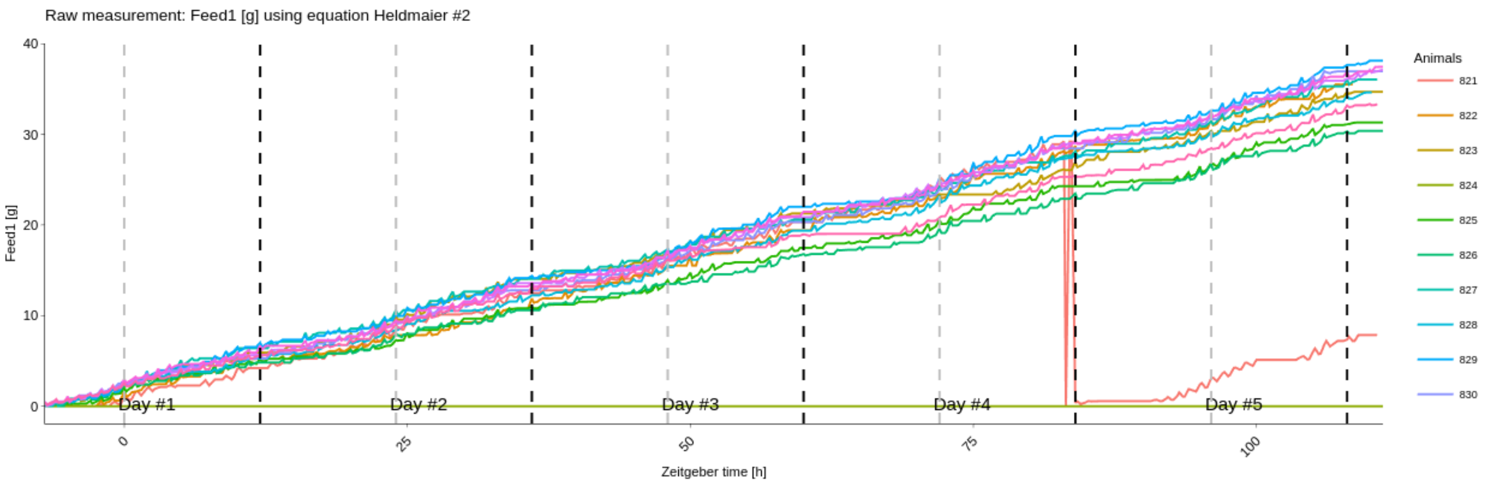


Drinking periods. Cumulative amount of water intake per animal:

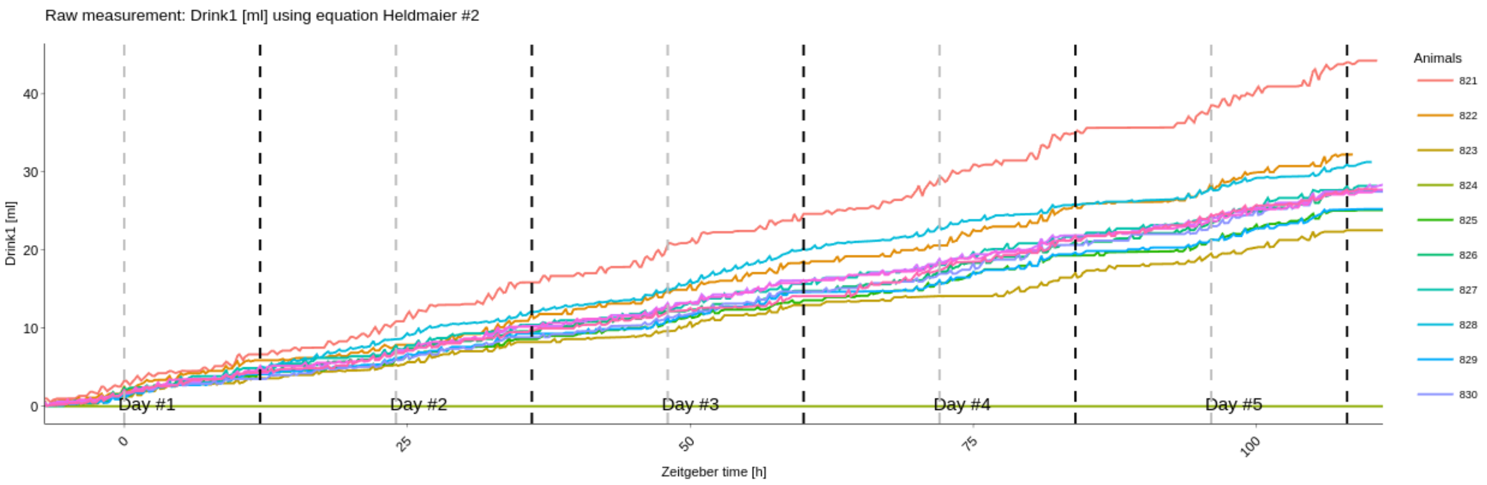

Body weight tracking. Current body weight (as determined by animals sporadically using the scale installed in the metabolic cage) over time per animal:

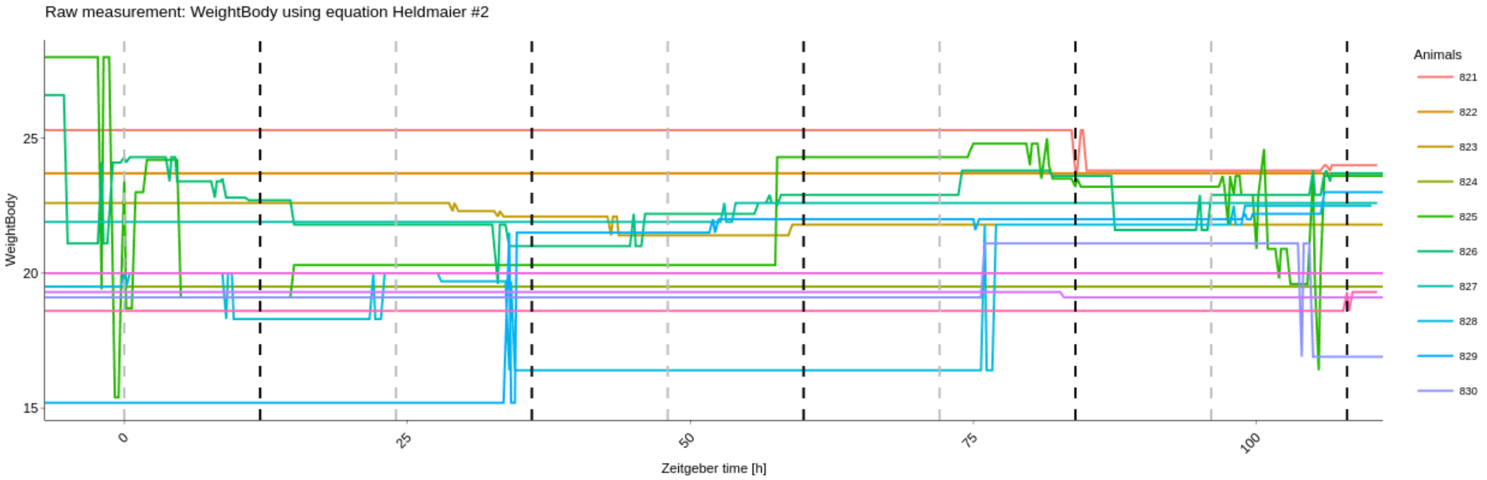

#### 6.3 III. Dataset generation from the IMPC database

The International Mouse Phenotyping Consortium (IMPC) contains raw data from a variety of indirect calorimetry experiments for different genotypes (and knockouts) as well as metabolic phenotypes.

When experimenters lack specific indirect calorimetry experiments, and still wish to analyze a knockout experiment for a given genotype (e.g. Ucp1 or Adipoq), the IMPC database might become useful.

While there exists a programmatic access through the Solr/Apache REST API to the IMPC database, complete indirect calorimetry datasets cannot be retrieved directly. However indirect calorimetry datasets can be manually created from separate datasets stored in the IMPC databases, tracking $O_2$, $CO_2$, etc. and additional metadata.

To allow Shiny-Calorie users to easily compile indirect calorimetry datasets for their choice of genotype from the IMPC database, a Python package called `calorimetry-tools`, cf. Listing 1. The package automatizes the generation of indirect calorimetry datasets from the IMPC database and stores the result in the TSE 6.3 row-based format which is directly loadable in Shiny-Calorie. The Python package can also be called via `reticulate` from GNU/R directly from Shiny-Calorie, however a pre-assembly of indirect calorimetry datasets is recommended, since large data retrieval from the IMPC database will take some processing time.

```python
from calorimetry_tools import *

def convert_from_impc_to_tse_file(gene_symbol):
    """Wrapper to convert from IMPC dataset to a TSE file"""
    # retrieve calorimetry measurements for a given gene symbol
    measurements = get_measurements_for_gene_symbol(gene_symbol)

    # combine these calorimetry measurements and store as data frame
    df = combine_measurements_for_gene_symbol(gene_symbol, measurements)

    # convert and write df to TSE PhenoMaster v6 row format
    write_tse(df, gene_symbol, f"results/TSE_file_for_{gene_symbol}.csv")

if __name__ == "__main__":
    # Example gene symbols
    gene_symbols = ["Ucp1", "Adipoq"]

    for gene in gene_symbols:
        convert_from_impc_to_tse_file(gene)
```

Listing 1: Example code to generate indirect calorimetry datasets for gene symbols Ucp1 and Adipoq from the IMPC database. The `calorimetry-tools` package makes internally use of the Apache/Solr REST API provided by the IMPC. Package is available on Github: `https://github.com/stephanmg/calorimetry-tools`.

#### 6.4 IV: Wavelet analysis to study ultradian rhythms

When high-frequency measurements ($< 60$ secs measurement interval) are available, the study of ultradian rhythms (URs) becomes possible. The study of URs has many important implications for unraveling the underlying cellular mechanisms of the complex behavior of for instance Djungarian hamsters.

Shiny-Calorie allows to study high-frequency recordings from the TSE Calobox indirect calorimetry system (single chamber) and to conduct wavelet analysis to detect URs, cf. Figs. S4-S5. Depicted is the power spectrum and the major significant frequency components based on the oxygen consumption which is proportional to the metabolic rate. Shiny-Calorie allows all calculated, derived and raw quantities to be analyzed by wavelet analysis. Interpolation to regularly-spaced time grids, pre-smoothing with LOESS (locally estimated scatterplot smoothing) or the Savitzy-Golay filter to add a correction for the delay of the measured gas exchange signals (properties of the measurement chamber) through z-transformation [10] (instantaneous metabolic rate) are subject to configuration by users, Shiny-Calorie provides reasonable defaults. Significant periods are marked in the periodogram with red circular markers.

### References

[1] L. Seep, S. Grein, I. Splichalova, et al. From Planning Stage To FAIR Data: A Practical Metadatasheet For Biomedical Scientists. *Sci Data*, 11, 2024.

[2] Patrick C. Even and Nachiket A. Nadkarni. Indirect calorimetry in laboratory mice and rats: principles, practical considerations, interpretation and perspectives. *American Journal of Physiology-Regulatory, Integrative and Comparative Physiology*, 303(5):R459–R476, 2012. doi: 10.1152/ajpregu.00137.2012. URL `https://doi.org/10.1152/ajpregu.00137.2012`. PMID: 22718809.

[3] G. Heldmaier and S. Steinlechner. Seasonal pattern and energetics of short daily torpor in the Djungarian hamster, Phodopus sungorus. *Oecologia*, 48:265—270, 1981. doi: https://doi.org/10.1007/BF00347975.

[4] J. B. de V. Weir. New methods for calculating metabolic rate with special reference to protein metabolism. *The Journal of Physiology*, 109(1-2):1–9, 1949. doi: https://doi.org/10.1113/jphysiol.1949.sp004363. URL `https://physoc.onlinelibrary.wiley.com/doi/abs/10.1113/jphysiol.1949.sp004363`.

[5] E. Ferrannini. The theoretical bases of indirect calorimetry: A review. *Metabolism*, 37(3):287–301, 1988. ISSN 0026-0495. doi: https://doi.org/10.1016/0026-0495(88)90110-2. URL `https://www.sciencedirect.com/science/article/pii/0026049588901102`.

[6] G. Lusk. *The Elements of the Science of Nutrition*. Sanders, Philadelphia, PA, 1928.

[7] M. Elia and G. Livesey. Energy Expenditure and Fuel Selection in Biological Systems: The Theory and Practice of Calculations Based on Indirect Calorimetry and Tracer Methods. In *Metabolic Control of Eating, Energy Expenditure and the Bioenergetics of Obesity*. S.Karger AG, 09 1992. ISBN 978-3-8055-5595-1. doi: 10.1159/000421672. URL `https://doi.org/10.1159/000421672`.

[8] E. Brouwer. Report of sub-committee on constant and factors. *Energy metabolism*, 11:441–443, 1965.

[9] Martin Klingenspor, Tobias Fromme, and Stefanie Maurer. Ablation of uncoupling protein 1 causes opposing changes of glucose uptake in brown and brite fat compatible with regulation of Glut 4 gene expression. *The FASEB Journal*, 34(S1):1–1, 2020. doi: https://doi.org/10.1096/fasebj.2020.34.s1.06713. URL `https://faseb.onlinelibrary.wiley.com/doi/abs/10.1096/fasebj.2020.34.s1.06713`.

[10] G. Bartholomew, D. Vleck, and C.M. Vleck. Instantaneous measurements of oxygen consumption during pre-flight warm-up and post-flight cooling in sphingid and saturniid moths. *J. Exp. Biol.*, pages 17–32, 1981.
